# Cell shape changes during larval body plan development in *Clytia hemisphaerica*

**DOI:** 10.1101/864223

**Authors:** Yulia Kraus, Sandra Chevalier, Evelyn Houliston

## Abstract

The cnidarian “planula” larva shows radial symmetry around a polarized, oral-aboral, body axis and comprises two epithelia cell layers, ectodermal and endodermal. This simple body plan is set up during gastrulation, a process which proceeds by a variety of modes amongst the diverse cnidarian species. In the hydrozoan laboratory model *Clytia hemisphaerica,* gastrulation involves a process termed unipolar cell ingression, in which the endoderm derives from mass ingression of individual cells via a process of epithelial-mesenchymal transition (EMT) around the future oral pole of an epithelial embryo. This contrasts markedly from the gastrulation mode in the anthozoan cnidarian *Nematostella vectensis*, in which endoderm formation primarily relies on cell sheet invagination. To understand the cellular basis of gastrulation in *Clytia* we have characterized in detail successive cell morphology changes during planula formation by Scanning and Transmission Electron Microscopy combined with confocal imaging. These changes successively accompany epithialization of the blastoderm, EMT occurring in the oral domain through the bottle cell formation and ingression, cohesive migration and intercalation of ingressed cells with mesenchymal morphology, and their epithelialization to form the endoderm. From our data, we have reconstructed the cascade of morphogenetic events leading to the formation of planula larva. We also matched the domains of cell morphology changes to the expression of selected regulatory and marker genes expressed during gastrulation. We propose that cell ingression in *Clytia* not only provides the endoderm, but generates internal forces that shape the embryo in the course of gastrulation. These observations help build a more complete understanding of the cellular basis of morphogenesis and of the evolutionary plasticity of cnidarian gastrulation modes.

## Introduction

Dynamic changes in the shape of individual cells underlie tissue remodeling during many of the morphogenetic events that sculpt the body of metazoan embryos, notably including the key event of gastrulation during which the germ layers and the embryo axes are set up (Solnica-Krezel & Sepich, 2012). Two morphogenetic processes driven by individual cell shape changes are very common during gastrulation: bending of cell sheets (e.g. invagination) and cell detachment from epithelia in a process of epithelial-mesenchyme transition (EMT) (Pearl et al., 2017; Shook & Keller, 2003). Both these processes involve transformation of columnar epithelial cells into a wedge shape by constriction of the apical cortex (Sawyer et al., 2010). At the cellular level such shape changes are driven by regionalized cortical enrichment and activation of myosin (Mason et al., 2013; Sherrard et al., 2010), as well as modulation of cell adhesion molecules and junction proteins to control interactions between neighbors (Buckley et al., 2014; Lecuit & Yap, 2015). Embryonic transcription programs combined with biomechanical feedback mechanisms guide changes in this cellular machinery through embryogenesis, with expression of key transcription factors such as Brachyury, Snail and Beta-catenin both directing and responding to these changes (Brunet et al., 2013; Pukhlyakova et al., 2018).

While much attention has focused on understanding individual morphogenetic processes, much remains to be understood about how these processes coordinate with each other or act ‘globally’ to shape the embryo (Beloussov, 2015; Miller & Davidson, 2013; Williams & Solnica-Krezel, 2017). This issue of global morphological integration has so far been addressed in animal models covering a quite restricted phylogenetic range such as nematode (Marston et al., 2016)*, Drosophila* (Rauzi et al., 2015), amphibians and birds (Firmino et al., 2016; Keller & Winklbauer, 1992; Shook et al., 2018), and ascidians (Sherrard et al., 2010). Much broader sampling will be needed to understand the various mechanisms that integrate morphogenetic movements of embryonic cells, and how this contributes to the evolution of gastrulation modes within the animal kingdom (Stower and Bertocchini, 2017).

Studies in Cnidaria can be very informative for understanding the cellular bases of developmental mechanisms and other morphogenetic processes. They can provide insights not only through comparisons with more traditional experimental model species from Bilateria, but also through comparisons between cnidarian species, which demonstrate extreme evolutionary and developmental plasticity. This is illustrated nicely by gastrulation, where cnidarians species deploy widely differing modes and cell behaviors to create the cnidarian planula larva (Kraus & Markov, 2017). Cnidaria planulae have a simple diploblastic organization with just two epithelial layers, endoderm and ectoderm organized around a polarized oral-aboral body axis. In some species the inner layer (endoderm) forms principally by cell sheet invagination from a hollow blastula, as described in scyphozoans (Berrill, 1949) and anthozoans (sea anemones, some corals: Gemmill & MacBride, 1920; Kraus & Technau, 2006; Magie et al., 2007; Marlow & Martindale 2007). In marked contrast, some hydrozoans and some anthozoans (octocorallians) develop via a solid morula stage, and gastrulation proceeds via germ layer delamination (hydrozoans: Harm, 1903; Muller-Cale, 1914, Kraus et al., 2014; Burmistrova et al., 2018; anthozoans: Matthews, 1917; Dahan & Benayahu, 1998). Endoderm can also form exclusively by ingression of individual cells from the future oral pole of the blastoderm, as is the case in *Clytia (=Phialidium)* species (Metchnikoff, 1886; Byrum, 2001).

We characterize in detail here cell morphology during embryonic development in *Clytia hemisphaerica. Clytia* exhibits the ‘typical’ complex life cycle of the class Hydrozoa: The adult stage (medusa) buds asexually from an ‘immortal’ juvenile stage (polyp colony). The medusa produces gametes daily for external fertilization and the resulting embryo develops into a planula larva. The planula metamorphoses after a few days into a sessile primary polyp, the founder of a new colony (Leclère et al., 2016). *Clytia* is a very useful experimental model for cell and developmental biology as well as evolutionary comparisons, being largely transparent and easy to manipulate in the laboratory. Inbred strains, genomic tools and knockdown techniques are now available (Leclère et al., 2019). The first studies of its embryonic development date back to the nineteenth century, when the mode of gastrulation by ingression of individual presumptive endoderm cells from one pole of the blastula was first described (Metchnikoff, 1886*; C. flavidula* = *C. hemisphaerica*). Using *C. gregarium,* the site of this “unipolar cell ingression” was later demonstrated experimentally to correspond to the animal pole of the egg (Freeman, 1981), and cell ingression was followed by marking experiments and morphometric analysis (Byrum, 2001). Molecular studies in *C. hemisphaerica* have shown that cell ingression occurs from specialized oral territory established in the blastula by the action of maternally localized mRNAs, which trigger Wnt pathway activation (Momose et al., 2008; Momose & Houliston, 2007). Genes activated downstream of Wnt/beta-catenin signaling in this oral domain include orthologs of known gastrulation regulators, notably two paralogous brachyury genes *CheBra1* and *CheBra2* (Lapébie et al., 2014).

Using *in vivo* imaging, confocal laser scanning microscopy (CLSM), Scanning Electron Microscopy (SEM) and Transmission Electron Microscopy (TEM), we describe here the succession of shape changes for cells of the *Clytia* embryo from cleavage until the planula stage, in relation to their position within the embryo and to the expression domains of selected genes. The characterization of cell shapes in relation to the overall morphology of the embryo during gastrulation allows us to reconstruct the morphogenetic events leading to the formation of planula larva and to predict the forces and tensions generated by the regionalized cell shape changes, ingressions, migrations and intercalations that characterize this particular gastrulation mode.

## Materials and Methods

### Animals and embryo cultures

*Clytia* embryos were obtained from laboratory Z strain adults (Leclère et al., 2019). Gametes were collected following light-induced spawning of adult medusae, mixed within one hour to achieve fertilization, and cleaving embryos collected into millipore filtered Red Sea Salt brand artificial sea water (MFSW) for culture. Embryonic development proceeds reliably at temperatures between 17- 21°C. Unless otherwise stated we express times of developmental events as hours post fertilization (hpf) at 18°C. Time lapse movies were made using Zeiss Axiophot-100 or Zeiss AxioObserver inverted microscopes. DIC still images were taken on an Olympus BX51 microscope. For early stages embryos were mounted in a small drop of MFSW between two coverslips separated by a moist spacer of filter paper and sealed with silicone grease. At the early gastrula stage the embryos start to swim by beating of ectodermal cilia, so they were embedded in 1.0 or 1.5% low melting point agarose (BioRad) in MFSW in a glass-bottomed petri dish to keep them in the field of view. Note that this does not prevent ciliary beating; the embryos continue to spin within an agar capsule. FIJI (https://imagej.net/Fiji) was used to adjust the light level across images stacks (‘bleach correction’ function), and to convert movies to .avi format, as well as for measurements of embryo dimensions.

### Scanning and transmission electron microscopy (SEM and TEM)

Embryos were fixed overnight at room temperature in the following fixative: one volume of 2.5% glutaraldehyde, four volumes of 0.4 M Na-cacodylate buffer (pH 7,2), and five volumes of microfiltered seawater (1,120 mOsm) (Ereskovsky et al., 2007), and then were transferred into 1.25% glutaraldehyde / 0.2M Na-cacodylate buffer (pH 7.2) and stored at +4°C. Before further processing, samples were post-fixed in 1% OsO4 / 0,2M Na-cacodylate buffer (pH 7,2) for 1 h and washed with the same buffer. Further processing was performed as described in Fritzenwanker et al. (2007). Samples for SEM were examined by the CamScan S-2 and JSM-6380LA scanning electron microscopes. Ultrathin sections were examined by the JEM-1011 transmission electron microscope with a Gatan ES500W Model 782 camera. Electron microscopy was performed at the Electron Microscopy Laboratory (Shared Facilities Centre of M.V. Lomonosov Moscow State University).

### Fluorescence staining and Confocal Microscopy

For staining of nuclei and cell contours, embryos were fixed for 2 hours in 4% formaldehyde in Immunobuffer (0.1M HEPES pH 6.9, 50mM EGTA, 0mM MgSO4, 80mM Maltose, 0.2% Triton-X-100) for two hours at room temperature. Following washes in PBS containing 0.01% Triton X100 (PBStx), actin cortices were staining by overnight incubation at 4°C in 1:100 rhodamine-phalloidin (stock in methanol; Fisher Scientific #10125092). For TUNEL staining using the “In Situ Cell Death Detection Kit TMR” (Roche #12156792910), embryos were fixed for four hours in Immunobuffer. Samples were then successively extracted/permeabilized by treatment with methanol (4°C for 2 hours to several days); 0.2% Triton X-100 in 0. 1% Sodium Citrate pH6.0 (30 minutes at room temperature): Proteinase K (1/25 of stock solution from the Click-iT™ Plus TUNEL assay kit - Molecular Probes #C10617 for 30 minutes at room temperature). Between each treatment, embryos were washed two times or more in PBStx. TUNEL labelling was performed in 1.5 ml eppendorf tubes containing groups of about 30 embryos in 30-40 µl PBS to which was added 84µl of reaction mix comprising 72µl Labelling solution and 12µl Enzyme solution from the Roche kit. After mixing, tubes were incubated for 2 hours at 37°C. For all fluorescence-stained embryos, PBStx washes and staining of DNA with 1 µg/ml Hoechst 33258 (Sigma-Aldrich #94403) preceded mounting in Citifluor AF-1 antifade mountant. All **s**amples were imaged on a Leica SP5 confocal microscope.

### In situ hybridization

Single in situ hybridization was performed using DIG labelled antisense probes exactly as described in Lapébie et al (2014). Double in situ hybridization used fluorescein labelled probe for *CheBra1* and a DIG labelled probe for *CheSnail,* detected using Peroxidase and Alkaline phosphatase conjugated antibodies with NBT or Fast red substrates respectively were performed as in (Jager et al., 2011). All genes and probes used here were described in Lapébie et al (2014) except *CheSnail* (accession number MN657238), which we isolated from our EST collection (Chevalier et al., 2006), and *Sox10* (Jager et al., 2011).

## Results

### Overview of *Clytia* normal development

The main events in the course of *Clytia* planula larva formation are cleavage, blastocoel formation, blastoderm epithelization, compaction, oral cell ingression, aboral-wards endodermal cell migration within the blastocoel, gastrula elongation and epithelization of the endoderm (see also outline in the sister study: van der Sande et al., 2020). These events are illustrated by light micrographs of the highly transparent embryo in Figures 1 and 2 and in the time-lapse movies of Supplementary Movies M1 and M2 (with stills provided in the corresponding Supplementary Figures S1 and S2). Cleavage divisions are holoblastic and unipolar, being initiated by nuclei/spindles positioned in the peripheral yolk-poor layer. The first cleavage division occurs about 50 minutes after fertilization and initiates at the animal/oral pole of the zygote, progressing to the aboral pole (Fig. 1A, B). Each subsequent cleavage cycle takes about 30 minutes. Starting from the second cleavage, blastomeres form lamella-like cytoplasmic projections and interact with their non-sister neighbors, thus pulling/crawling against each other (Fig. 1C, C’, D). This results in “twisting” of the forming blastomeres relative to each other and compact cell packing, which morphologically resembles spiralian cleavage (Fig. 1E, F),but is achieved by a distinct mechanism. At the 32-cell stage a blastocoel appears (Fig. 1H) defining the blastula stage. The blastocoel expands during the next rounds of synchronous divisions as illustrated at the 128-cell stage in Fig. 1I. Unlike in some other cnidarian species, cleavage division in *Clytia* embryos are always complete and are highly synchronous until the early blastula stage. These cleavage divisions are symmetric, but since the interactions between newly formed blastomeres are quite loose the initial shape of the embryos is variable and irregular At about 7 hpf as epithelialization commences and individual cells flatten against each other the blastula acquires a more spherical shape.

**Figure 1.**
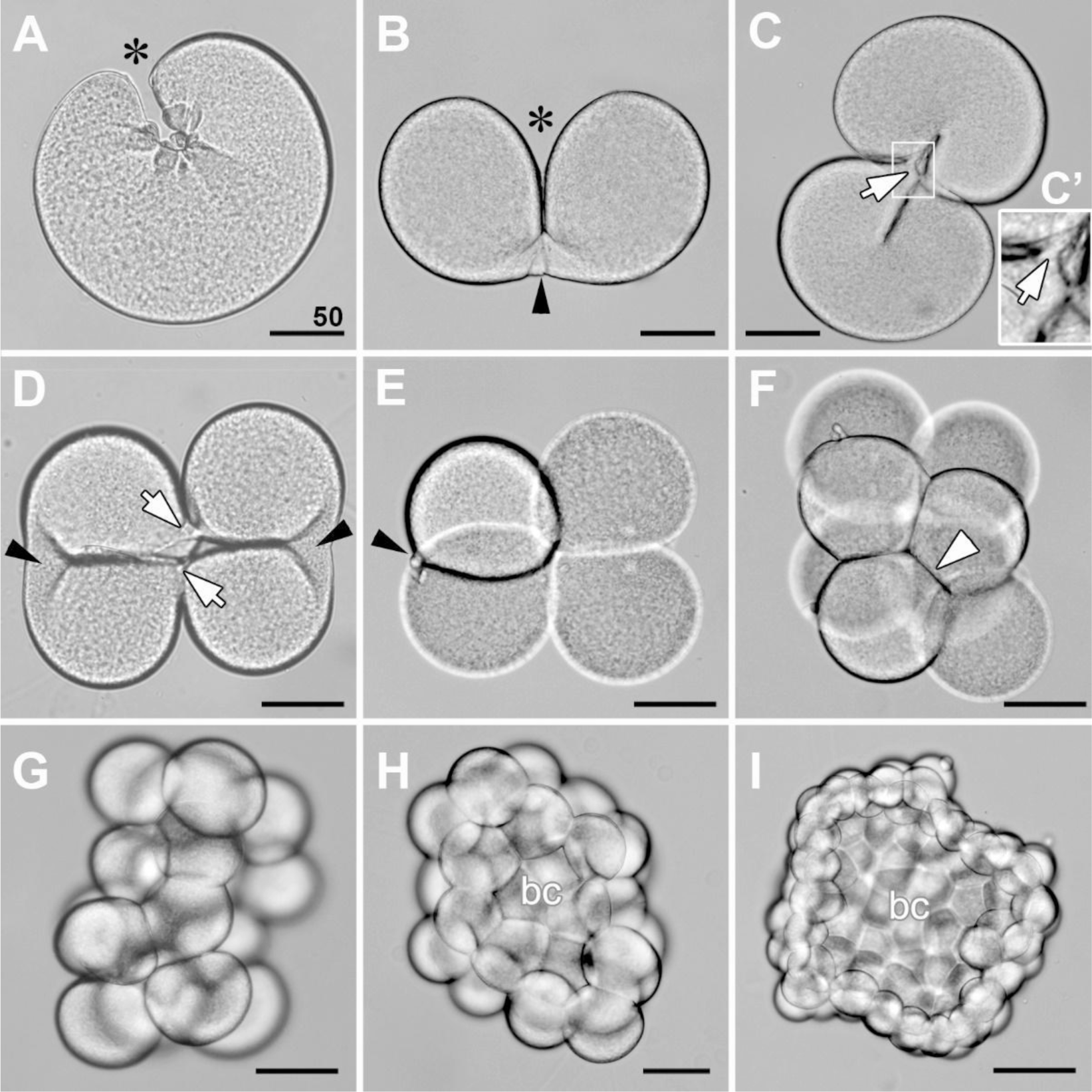
Cleavage divisions to form the early blastula. Light microscopy images of living embryos. (A) Initiation of the first cleavage furrow: asterisk marks the site of furrow initiation, corresponding to the egg animal pole. (B) Elongation of the first cleavage furrow. Black arrowhead points to the cytoplasmic bridge between the forming blastomeres. (C, D, E) Transition between two-cell stage and four-cell stage, viewed from the animal pole. Two second cleavage furrows start to form in (C); (C’) is enlarged from the area framed in (C). White arrows mark lamellae connecting the non-sister blastomeres (C, C’). Forming blastomeres often twist against each other as the furrow advances, resulting in tetrahedral configurations as in (E). (F) Eight-cell stage embryo with blastomere packing in this example superficially reminiscent of spiral cleavage. Arrowheads point to contacts between non-sister blastomeres. (G) 16-cell stage. (H) 32-cell stage, appearance of the blastocoel (bc). (I) 128-cell stage. Cleavage divisions in *Clytia* are very synchronous- see Movies S1 and S2. Note that in (G-I-hat blastomeres are in focus. All scale bars - 50 µm.

**Figure 2.**
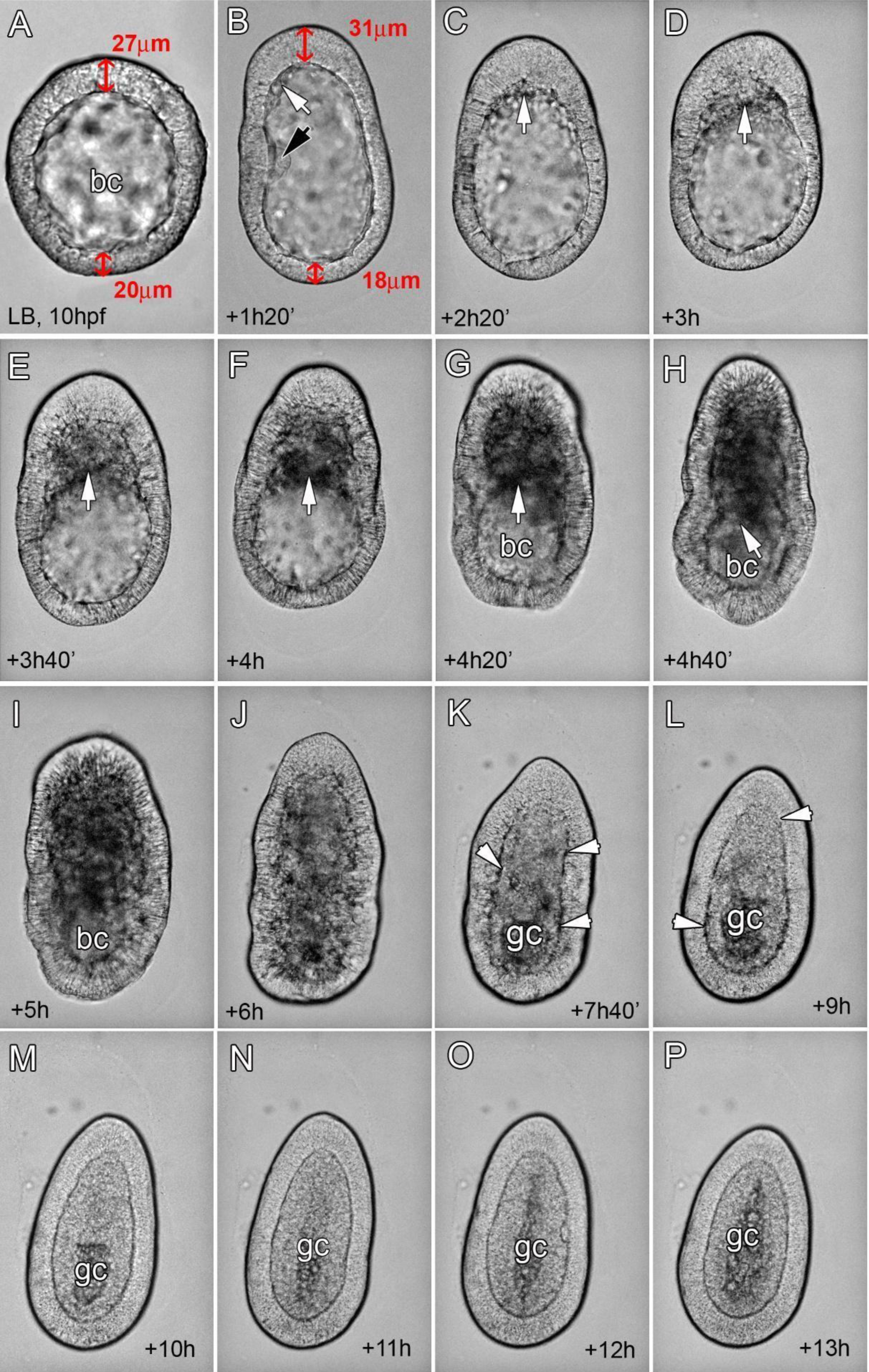
Development from the late blastula to the planula stage. Light microscopy images of live embryos; oral pole at the top in all cases. (A) Embryo at the late blastula stage; thickening of the oral half blastoderm compared to the aboral blastoderm, indicated by red double-headed arrows, is the first morphological sign of the OA axis. (B - P) Frames taken from a time-lapse movie of an individual gastrulating embryo selected from a culture at 18°C and embedded in low-melting-point agarose. The time from the start of the recording is indicated in each frame -note that development from this time proceeded at a higher temperature (about 20°C). Another consequence of the embedding is the exaggerated elongation along the OA axis at the onset of gastrulation - compare with unembedded embryos in Fig. S3 (B) Early gastrula stage, at the very beginning of gastrulation. White arrow points to the first ingressing cells. Black arrow points to a fold in the blastoderm which arose during cleavage. (C - I) Successive stages of gastrulation. White arrows point to the front of the mass of ingressed cells moving towards the aboral pole. Note a gradual reduction of the blastocoel (bc). (J) Parenchymula stage. Gastrulation is complete; blastocoel is filled with ingressed cells. (K - P). Development of the planula larva. White arrowheads point to the ECM layer separating the germ layers. A gastrocoel (gc) starts to form in the aboral endoderm and spreads towards the oral pole as ingressed cells reorganize to form a gastrodermal epithelium. The morphology of the oral/posterior pole is pointed and of the aboral/anterior pole is rounded. Scale bar 50 µm

During the blastula stage, the diameter of the embryo becomes smaller due to a “compaction” process accompanying epithelialization. Measurement of the cross sectional areas of embryos from two batches cultured at 17°C-18°C showed reductions of 11.6 +/-7.0% and 8.8 +/-7.2% respectively (details in Supplementary Fig. S3). At the late blastula stage the diameter of the embryo becomes smaller due to a compaction process accompanying epithelialization, and then regional differences in the thickness of the blastoderm are first observed (Fig. 2A and Supplementary Fig. S3). Embryos start to swim by beating of cilia on the blastoderm cells, oriented by a conserved planar cell polarity (PCP) mechanism operating along the oral-aboral axis (Momose et al. 2012). The onset of gastrulation is then marked by more pronounced local thickening of the blastoderm, marking the site of cell ingression and future oral pole of the embryo (Fig. 2B, C). As gastrulation progresses, the mass of ingressed cells gradually fills the blastocoel and the front of the mass displaces towards the aboral pole (Fig. 2D - I). This is accompanied by a characteristic change in overall embryo morphology: the oral pole of the gastrula becomes pointed, and the region that contains ingressed cells narrows compared to the more aboral region where the blastocoel is still empty. Around 6 hours after the onset of gastrulation (corresponding to roughly 20 hpf at 18°C), ingression is complete and the blastocoel filled with cells, defining the parenchymula stage (Fig. 2J). Over the next 24 hours gastrulation is completed by reorganization of the ingressed cells to form a new, distinct epithelial endodermal layer. A gastrocoel forms in the central core and a distinct extracellular layer develops, separating the differentiating ectodermal and endodermal epithelia (Fig. 2K - P; this process is characterized in more detail below as basal lamina formation). Both the formation of the gastrocoel and the separation of the ectodermal layers initiate at the aboral pole and progress towards the oral pole (compare images Fig. 2K - O). Differentiation of specific cell types then proceeds and the planula becomes competent to settle and metamorphose at about 72 hpf.

### Cellular morphology changes accompanying gastrulation

To understand the changes in cellular cell shape that underlie planula morphogenesis, we examined in detail embryos fixed at successive stages by a combination of Scanning Electron Microscopy (SEM), Transmission Electron Microscopy (TEM), and Confocal Laser Scanning Microscopy (CLSM) following staining of cell boundaries with fluorescent phalloidin and nuclei with Hoechst dye (Figs 3-8).

### Blastula stage

The major cellular event during the blastula stage is blastoderm epithelization. Early blastula embryos (3 - 6 hpf) have irregular shapes (Fig. 3A, B). Starting from the first cleavage stage and until 7hpf cell divisions are almost synchronous (Fig. 3C). At this stage the blastodermal cells do not show any morphological signs of apico-basal polarity, as they are rounded and their nuclei are positioned close to the cell center (Fig. 3B, D). Division synchrony is lost during the blastula stage (see Supplementary Movie M1). At 7-8 hpf the embryo reaches the mid-blastula stage, and its shape becomes more regular (Fig. 3E). Blastoderm cells show the first morphological signs of polarization along the apico-basal axis: cells start to adopt a columnar shape, their nuclei displacing towards the apical surface (Fig. 3E, E’, F). At the late blastula stage (9 - 10hpf) the embryo shape is almost spherical (Fig. 3G). The blastoderm cells show polarized ultrastructure: their apical domains are essentially free of yolk granules; the apical surface is covered with microvilli; the apical cortex is enriched with the cortical granules; electron dense intercellular junctions are localized in subapical area; yolk granules are displaced to the basal domain (Fig. 3H, H’). At this stage cell divisions clearly become asynchronous and all blastodermal cells are columnar (Fig. 3I). Well-developed cilia are present on the cell apical surfaces from approximately 8hpf (Fig. 3G,J). At about 10hpf blastodermal cells become more markedly columnar, with the apicobasal-axis: apical-width ratio reaching 2.6 +/- 0.16 (min 2.3; max 2.8 as measured on images of 10 cells from 5 embryos) (Fig. 3K - M), but the shape of individual cells is not uniform. In some cells the lateral surfaces are roughly parallel while in others the basal surfaces are either expanded or reduced relative to the apical surfaces (Fig. 3I, L, M).

**Figure 3.**
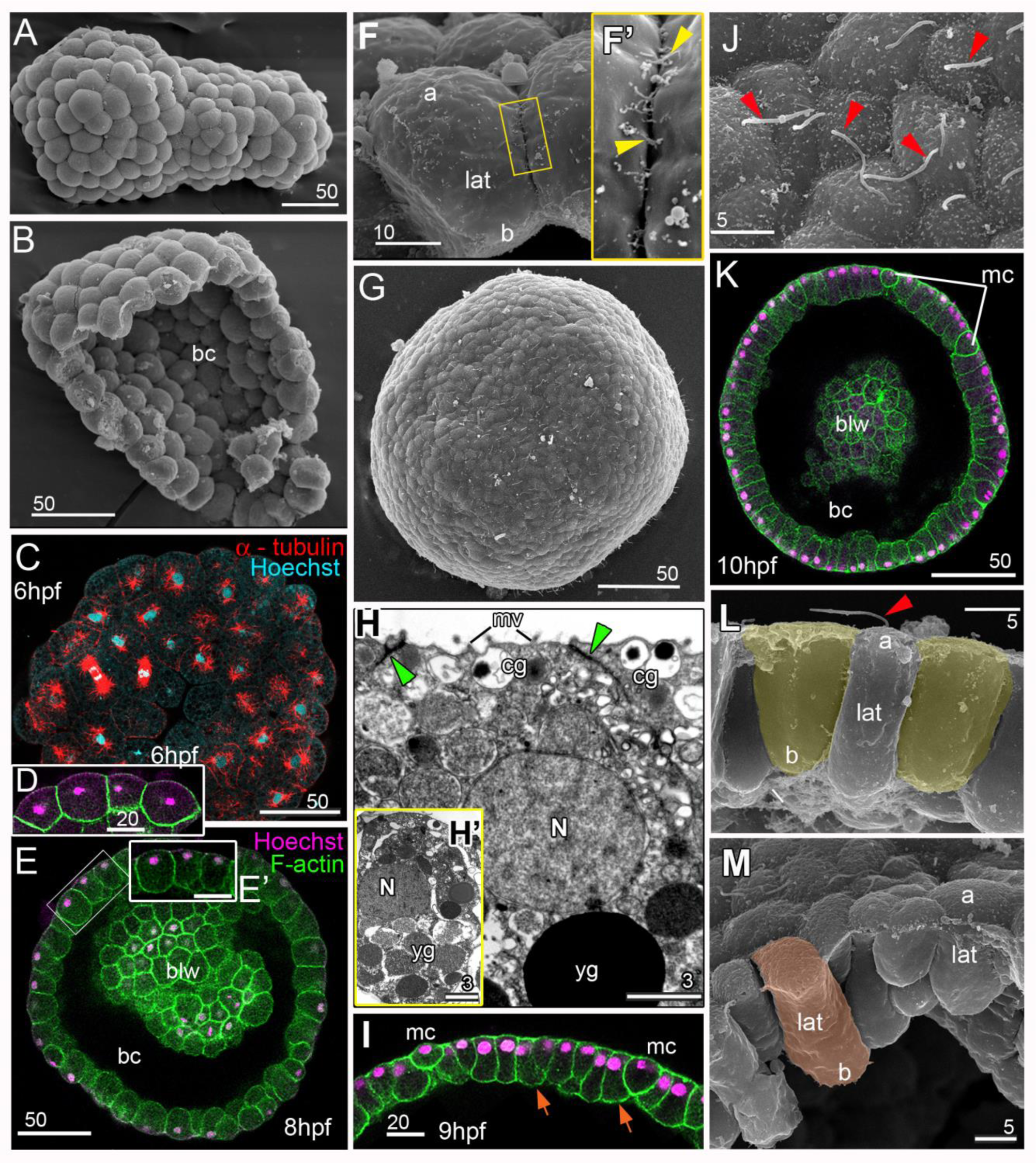
Gross morphology and cell morphology of blastula stage embryos. (A) SEM of an embryo at the early blastula stage (256 - 512 cells), with an irregular, non-spherical shape. (B) SEM of an early blastula split into two halves. Blastoderm cells have a rounded shape. (C) CLSM z-stack of an early blastula, the cells undergo synchronous division; mitotic spindles are labeled with anti-tubulin antibody (red) with Hoechst staining of DNA in cyan. (D) CLSM z-plane of early blastula cells. In this and in all other CLSM images, actin-rich cell cortices are labeled with phalloidin (green) and nuclei with Hoechst dye (magenta). The slightly stronger background apical Hoechst staining in panel D should not be confused with apical positioning of the nuclei. Cell shapes are irregular and nuclei central; bc - blastocoel. (E) CLSM of an embryo at the mid-blastula stage, blw - blastocoel wall. (E’) Cells framed in (E) are enlarged to the same magnification as (D) and (I) for direct comparison. The blastula shape has become more regular. Cells have elongated slightly along their apico-basal axes and nuclei have displaced to the apical pole of each cell. (F, F’) SEM of cells from a split mid-blastula stage embryo. (F’) Enlargement of the area framed in (F); short filopodia (yellow arrowheads) cover the lateral cell surface providing the cell-cell contacts; a - apical, lat - lateral, b - basal cell surfaces. (G-M) Late blastula stage: SEM (G, J, L, M), TEM (H, H’) and CLSM (I, K). (G) Surface view of an embryo, whose shape is close to spherical. (H) Apical and perinuclear areas of a blastoderm cell; cg - cortical granules, mv - apical microvilli, N - nucleus, yg - yolk granules, green arrows - sub-apical intercellular junctions. (H’) A cell of the same embryo showing basal localization of yolk granules and apical position of the nucleus. (I) CLSM z-plane of late blastula cells showing a variety of cell shapes and an asynchrony of cell divisions; magnification is the same (D) and (E’), orange arrows indicate cells with expanded basal domains; mc - rounded mitotic cells. (J) Apical surfaces of late blastula cells; each cell has a cilium (red arrowhead). (K - M) Images illustrating further elongation of blastoderm cells along the apico-basal axis and a variety of blastoderm cell shapes. A cell with a tapered basal end (left) and one with a cylindrical form (right) are highlighted in yellow in (L); cell with an expanded basal domain is highlighted in orange (M). Scale bars: A - E, G, K - 50µm; D, E’, I - 20 µm; F - 10 µm; H, H’ - 3 µm; J, L, M - 5µm.

The cell shapes characterized in the late blastula and in different regions of the embryos during successive stages of planula development, as described in the following sections, are summarized schematically in Figure 4.

**Figure 4.**
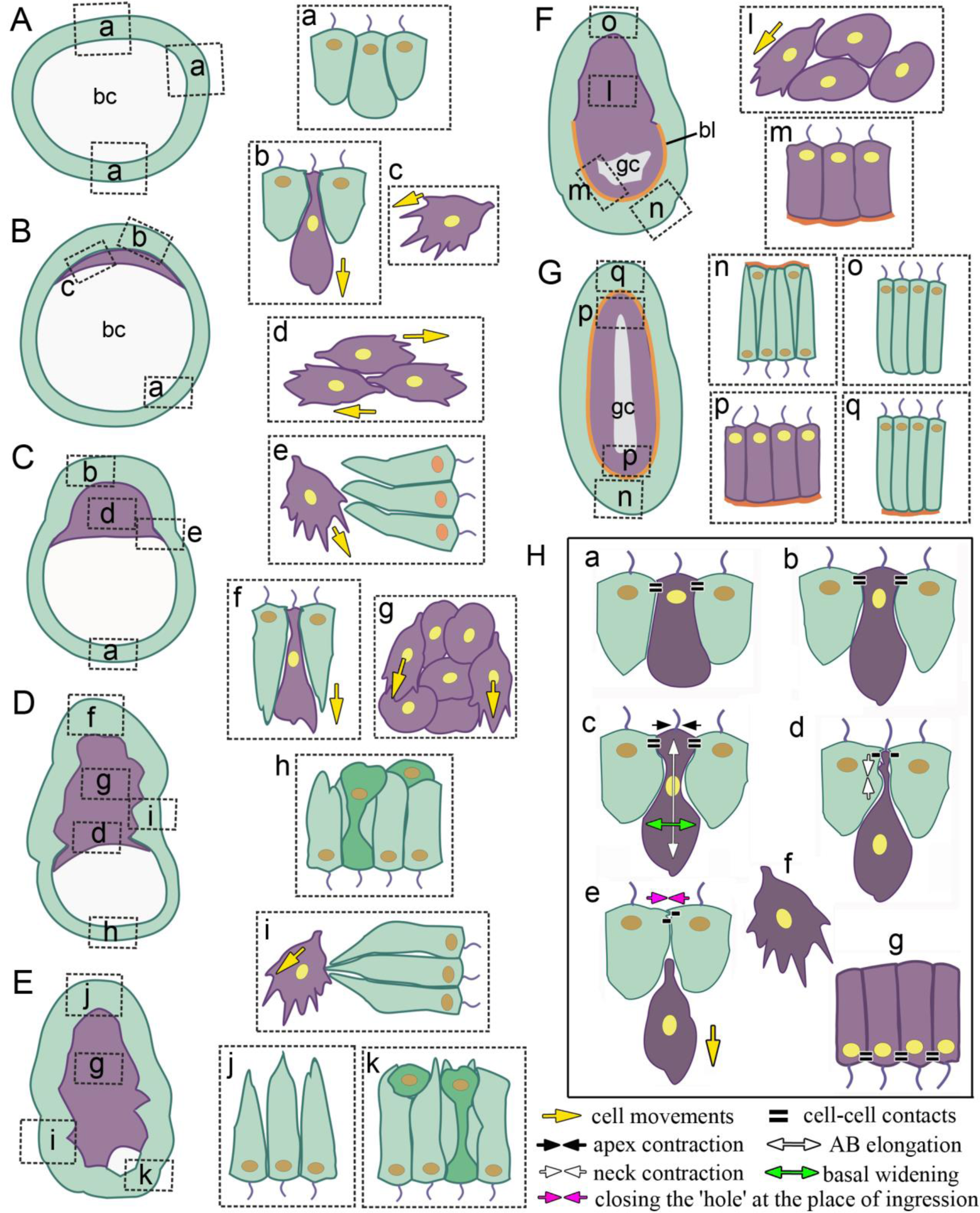
Scheme of cell shape changes during *Clytia* early development. The cartoons show the cell shapes (a - q) in different regions of embryos at successive stages of development (A - G). Embryos stages: (A) late blastula, (B) early gastrula, (C) mid-gastrula, (D) late gastrula, (E) late gastrula, during completion of blastocoel reduction, (F) parenchymula with gastrocoel formation starting, (G) 2 day planula. (H) The cartoons follow the changes in individual presumptive endoderm cells during: (a - d) successive stages of bottle cell formation, (e, f) cell ingression and migration (g) endoderm epithelization. Cells of the outer layer shown in green include both presumptive ectoderm and presumptive endoderm cells at early stages (A-D) but only ectoderm cells at later stages (E - G). Presumptive endoderm is in violet; basallamina/ECM (G, H) is in orange; yellow arrows show directions of movements of the future endodermal cells; black, white and pink arrows in H indicate specific cell morphology changes as indicated in the key; gastrocoel (F, G) is in light gray.

### Early gastrula stage - bottle cell formation

Starting around 9 hpf, the blastoderm becomes thicker on one side of the late blastula, marking the future position of the oral pole (Figs. 2A and 5A, B). The thickening of the oral pole blastoderm becomes pronounced at the onset of gastrulation (10 - 11 hpf; compare oral (top) and aboral(bottom) regions in Figs. 2B and 5A, B, B’). The oral domain of the late blastula - early gastrula is quite variable in size, covering about one third of the embryo. (The mean ratio of the oral blastoderm territory (see Fig. 5A) measured on sections as the contour between the most lateral bottle cells to the rest blastoderm was 2.3 ± 0.65; minimum 1.5; maximum 3.5; n = 18).

**Figure 5.**
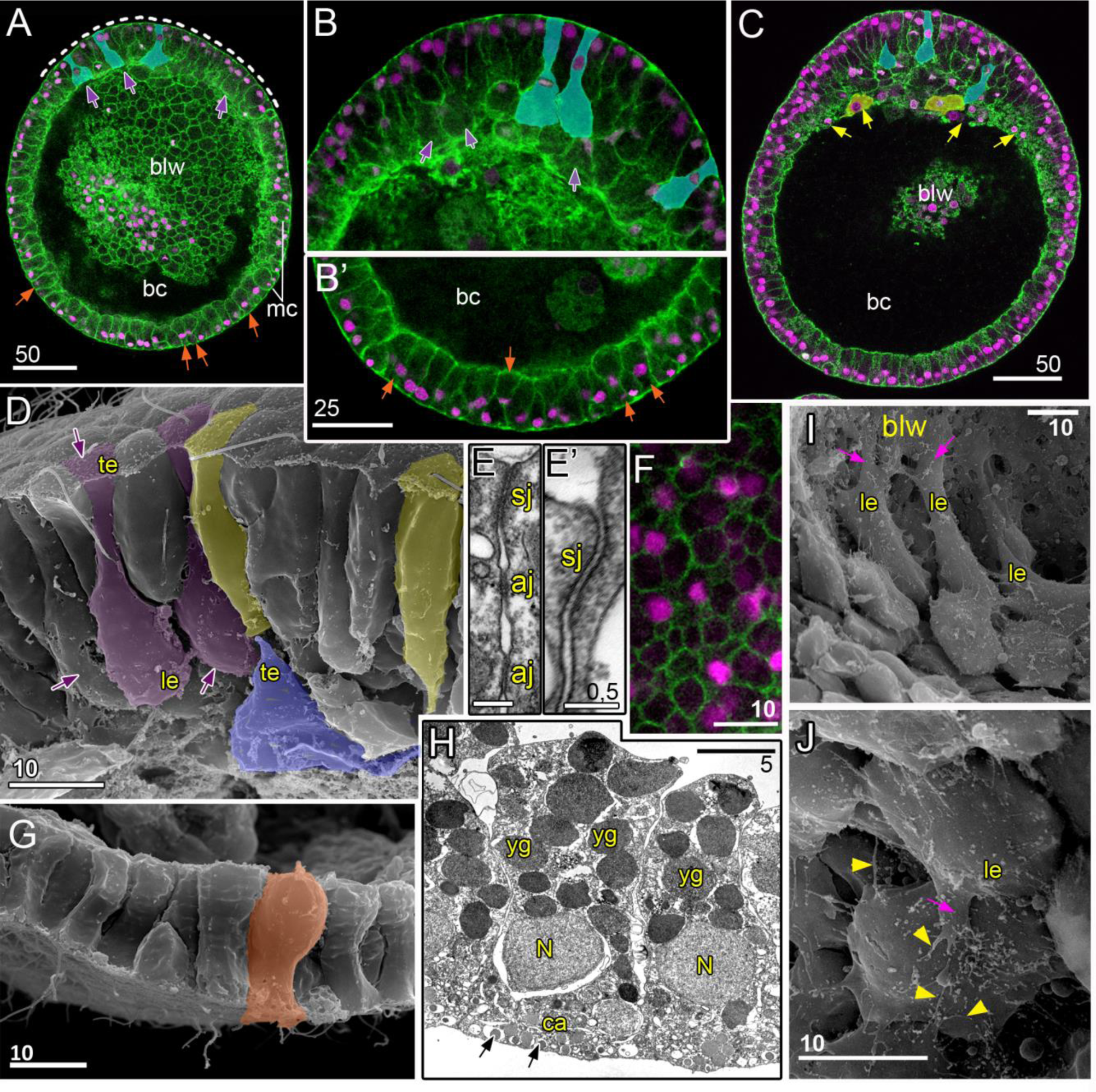
Early gastrula stage: distinct cell morphologies in the oral domain. (A, B, B’) CLSM images of embryos at the very beginning of gastrulation, before any cells have entered the blastocoel. The oral pole is towards the top in all images. In the oral blastoderm (A, B), violet arrows indicate bottle cells characterized by elongated apico-basal axes, contracted apical surfaces and expanded basal surfaces. Cells highlighted in turquoise are bottle cells at different stages leading to ingression, as indicated by the location of the nuclei in apical or central positions. These stages are characterized further in Figure 6. The white dotted line in (A) delineates the oral domain. In the aboral blastoderm (A, B’), orange arrows indicate cells with expanded basal areas, morphologically resembling bottle cells. (C) CLSM of an early gastrula embryo demonstrating ingression of oral cells has now commenced. Some ingressed cells (yellow arrows) have started to migrate inside the blastocoel. (D) SEM of oral cells of split embryos at an early gastrula stage. Oral bottle cells are highlighted in violet and a cell almost ingressed into the blastocoel in lilac. Columnar cells with relatively short apico-basal axes (“squat cells”) are highlighted in yellow; le - leading, te - trailing edges of bottle cells preparing to ingress. (E) TEM of sub-apical junctions between two oral cells; sj - septate junction, aj - adherence junction. (E’) Septate junction at higher magnification. (F) Confocal plane parallel and close to the apical surfaces of a field of aboral cells, showing a wide range of apical surface areas. (G) SEM of aboral cells of a split early gastrula, a cell with an expanded basal area is highlighted in orange. (H) TEM of aboral cells in an early gastrula; N - nucleus, yg - yolk granules, ca - cortical area, black arrows point to the cortical granules. (I) Ingressed cells migrating over the blastocoel wall. Their leading edges (le) are very pronounced and form wide lamellae (magenta arrows). (J) Ingressed cells located further from the blastocoel wall have small lamella and multiple filopodiae (yellow arrowheads). Scale bars: A, C -50 µm; B, B’ - 25 µm; D, F, G, I, J - 10 µm; E, E’ - 0,5 µm; H - 5 µm.

Examination of cellular morphology within the thickened oral area reveals a mixture of cell shapes, classifiable into two types of morphology (schematized in Fig. 4Bb). Cells of the first type (colored in yellow in Figs. 5 and 6) have short apico-basal axes and are columnar or wedge shaped. These are ordinary epithelial cells. Cells of the second type (colored in violet in Figs. 4, 5 and 6) have elongated apico-basal axes, expanded basal domains and frequently a reduced apical width. These features classically characterize ‘bottle cells’. We found bottle cells in *Clytia* to be distributed evenly within the oral territory, where ordinary epithelial cells are mixed with forming bottle cells (Figs. 5B, C, D and 6A’, D). All blastoderm cells, including forming bottle cells, were found to be connected to each other by subapical set of intercellular junctions, apical septate junctions and more basal adhesion junctions (Fig. 5E, E’).

**Figure 6.**
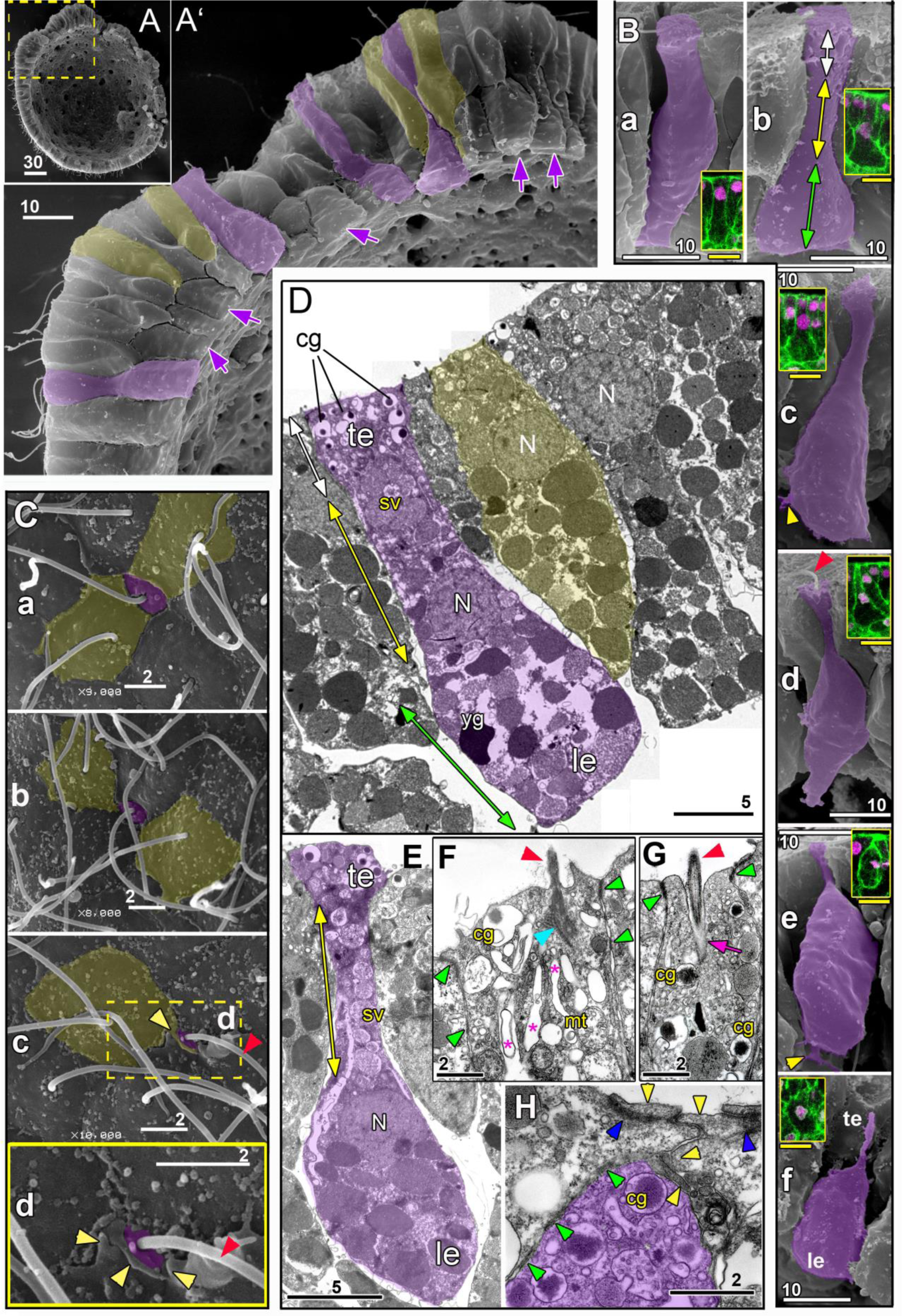
Morphology and ultrastructure of bottle cells at the early gastrula stage. (A) SEM image showing the cut surface of an embryo fixed and split at the early gastrula stage. (A’) Zoom on the oral area framed in (A). Multiple bottle cells (violet color, violet arrows) alternate with columnar cells with relatively short apico-basal axes (“squat cells” yellow color). (B) Successive stages (a - f) of bottle cell reshaping during ingression, SEM and CLSM images. (a) At the beginning, the cell apical domain (white double-headed arrow) is already slightly contracted and the nucleus is located apically. At the next step (b - c), the ‘neck’ region between the apical and basal areas (yellow double-headed arrow) becomes longer and narrower than the apical domain; the nucleus is displaced into the neck; the basal domain (green double-headed arrow) expands. (d) The apex contracts further, the cell neck becomes narrower and slightly contracts too. (e) The neck contracts further and the nucleus is positioned in the basal part of the cell. (f) The cell detaches from neighboring cells and ingresses into the blastocoel. Note the protrusions (yellow arrowheads) on the basal surfaces of bottle cells and cilium on the apical surface (red arrowhead). le - leading edge, te - trailing edge. (C) SEM of the oral surface of an embryo, successive stages of the ‘sinking’ of ingressing bottle cells (a - d); apical surfaces of bottle cells are highlighted in violet and of neighboring epithelial cells in yellow. Yellow arrowheads point to filopodia-like protrusions of epithelial cells extending over the surface of the sinking bottle cells; (d) zoom of the area framed in (c). (D) TEM of the early gastrula oral area: two forming bottle cells (one colored in violet) and two squat cells with short apico-basal axes (one colored in yellow). The bottle cell nucleus is positioned distant from the contracted apical domain, near the expanded basal domain in the ‘neck’ region between the apical and basal areas. Cortical granules (cg) are displaced deeper into the apical cytoplasm. N - nuclei; sv - putative storage vesicles; yg - yolk granules. (E) TEM of a bottle cell at a later stage of ingression than the cell shown in (D): the ‘neck’ is now narrower than the apical domain and the nucleus is displaced into the expanded basal area; storage vesicles are lined up along the cell neck. (F - H) TEMs of the apical domains of bottle cells at three stages of cell ingression. (F) At the beginning of ingression, bottle cell retains the cilium (red arrowhead, with blue arrowhead pointing to the cilium root) and sub-apical intercellular junctions (green arrowheads). Note the elongated shape of vacuoles (marked with stars) squeezed by the apical contraction; mt - mitochondria. (G) The next stage of apical contraction: the cilium deeps within a ‘pocket’ formed from the apical cell membrane (purple arrow); cortical granules are displaced from the apical surface; intercellular junctions are still preserved. (H) Apical domain of a cell that has already ingressed (violet color), covered by protrusions of neighboring epithelial cells closing the ‘hole’ at the place of ingression (yellow arrowheads). Well-developed intercellular junctions (blue arrowheads) are present between these protrusions. The ingressed cell retains sites of adhesion with neighboring epithelial cells (green arrowheads). Scale bars: A - 30µm, A’ - 10µm; B - 10µm (white scale bars - SEM images; yellow scale bars - CLSM images); C, F, G, H - 2µm; D, E - 5µm.

Across the rest of the embryo, corresponding to future aboral and lateral regions of the early gastrula, wedge-shaped or columnar cells also alternate with cells with expanded basal domains (Figs. 4B and 5A, B, G). As in the oral domain, the apical cell surface areas are variable (Fig. 5F), and many cells have expanded basal domains and reduced apical widths. The “basally expanded” cells in aboral/lateral regions did not show increased elongation along the apicobasal axis, distinguishing them from the true bottle cells observed in the oral domain. The nuclei of most aboral/lateral cells at this stage were located within the apical half of the cell, occupying surface or medial positions. TEM revealed that storage vesicles enriched in yolk are concentrated basal to the nucleus in all cells (Fig. 5H). These comparisons of cell morphology across the embryo indicate that local thickening of the oral blastoderm prior to gastrulation is caused by apico-basal elongation of a sub-population of cells undergoing bottle cell formation.

Within the oral domain, the widely variable morphologies of bottle cells (eg cells marked in turquoise in Fig. 5A, B, C and in violet in Fig. 6A’, see also cartoon in Fig. 4H) correspond to the stages of bottle cell formation described as a component of “primary” EMT, ie the EMT processes of early embryogenesis (Shook & Keller, 2003; see Discussion). Since EMT of these cells is not synchronized within the blastoderm, nearby cells are observed at different stages of the process (Fig. 6A’, B), finally detaching individually from the epithelial layer and adopting a mesenchymal morphology (Figs. 5D, I, J and 6Bf). Close examination of the cell apical surface revealed that epithelial cells surrounding the ingressing bottle cells form multiple filopodia-like protrusions covering the bottle cell apical surface (Fig. 6C). New junctional complexes are formed between the protrusions of the neighboring cells (Fig. 6H). In this way, the epithelial sheet remains sealed as the ingressing cells detach into the blastocoel.

From electron microscopy data, we could detail successive stages of bottle cell formation (Fig. 6B; schematized in Fig. 4H). First, the apical surface contracts slightly and the apico-basal starts to elongate (Fig. 6Ba). Concurrently, the basal part expands and a narrow neck forms, connecting distinct apical and the basal domains (Fig. 6Bb, D). At this stage the nucleus is located in the cell neck (Fig. 6Bb, D). At the next step, the apical surface of the bottle cell progressively reduces, and the neck becomes narrower (Fig. 6Bc, Bd). Constriction of the cell apical domain appears to force the cytoplasm and nucleus towards the basal domain (Fig. 6E -G). Cytoplasmic components become deformed along the apico-basal axis (such as the vacuoles in Fig. 6F) or align along the neck (such as the storage vesicles in Fig. 6E). The nucleus then displaces from the neck to the expanded basal part of the cell (Fig. 6E). Interestingly, ingressing cells retain their sub-apical junctions and cilia during most of the ingression process, even when the apical surface is reduced to a very small area (Fig. 6Bc, Cc, Cd, G). At the next step, the apex of bottle cell contracts further, while the neck region becomes narrower and slightly shorter (Fig. 6Bd, 6Be). Due to extreme contraction of the cell apex, cytoplasmic components located near the apical surface, such as the cortical granules in the Figs. 5H and 6D, move deeper, while sub-apical junctions are still retained in the bottle cells (Fig. 6G). Finally, the neck of the bottle cell contracts, and the cell leaves the epithelium (Figs. 6Bf, 6H; lilac cell in 5D).

In the course of ingression, each bottle cell gradually adopts morphological features of a migrating cell: its basal surface transforms into the leading edge, and its apical surface becomes the trailing edge upon detachment from the blastoderm (Fig. 6B). However, only few filopodial or lamellipodial protrusions are detected at the leading edge at this time (Fig. 6Bc, Be). These morphological observations suggest that bottle cells drop out of the oral blastoderm principally as a result of cell shape changes, and only start to migrate once in the blastocoel. Once ingressed, cells have a typical mesenchymal phenotype: they are fibroblastic in appearance with a flattened shape, leading edge – trailing edge polarity, numerous processes and invasive behavior (Hay, 2005); their morphology Indicates that they move actively by crawling on the blastocoel wall and neighboring cells including other ingressed cells (Figs. 5I, J; 7E, F, H). Cells migrating over the blastocoel wall develop very pronounced leading edges with wide lamella adorned with multiple filopodiae (Fig. 5I). These cells crawl towards the aboral pole of the embryo. Cells located further from the blastocoel wall appear to use other ingressed cells as a substrate for crawling. They form small lamellae and multiple filopodiae (Fig. 5J).

### Mid-gastrula stage

During the mid-gastrula stage (Fig. 7), embryos adopt a characteristic pear shape as the part of the embryo in which cells have entered the blastocoel becomes thinner. The embryo can thus be divided into two distinct parts, a rounded aboral part and a thinner, more pointed, oral part (Figs. 4C; 7A-D). The behavior of the ingressed cells during mid-gastrula stage appears highly coordinated. Flattened cells with mesenchymal morphology displace as streams or cohesive blocks in which cells are oriented end-to-end and side-to-side, aligning as layers perpendicular to the oral- aboral axis (Fig. 7C-E). The morphology and arrangement of these cells inside the blastocoel indicates that central cells migrate relative to each other and intercalate along the elongating oral- aboral axis (Figs. 4Cd; 7D, E, F, H), while those close to the blastocoel wall migrate along it towards the aboral pole (Fig. 7C, D, G, J, K). Migrating cells form multiple spot-like sites of adhesion between each other (Fig. 7I).

**Figure 7.**
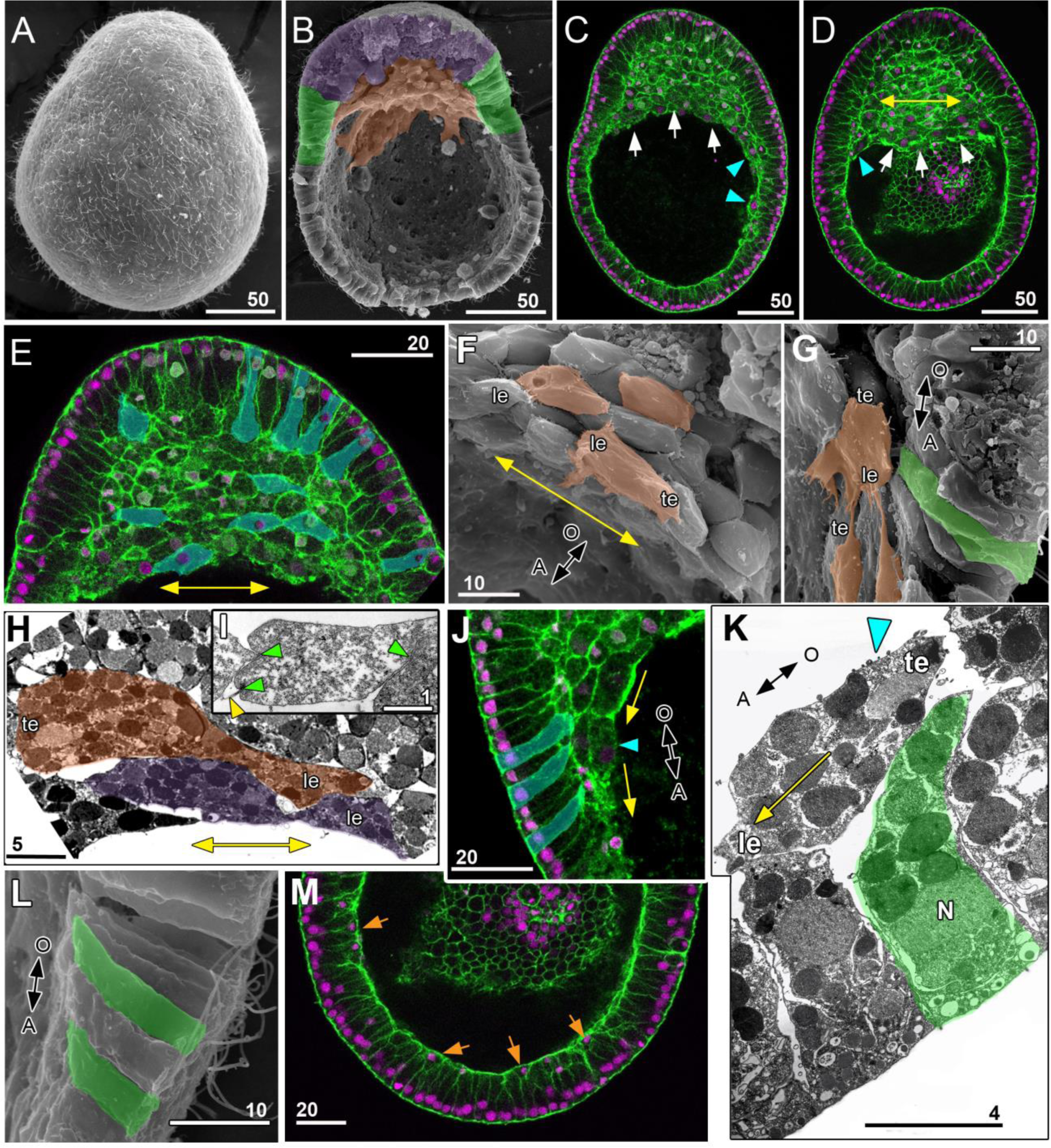
The mid-gastrula stage: continued cell ingression, migration of ingressed cells. (A - D) Mid-gastrula stage embryos positioned with their pointed oral poles towards the top in all panels. (A) SEM image, side view. (B, C, D) Successive stages of cell ingression illustrated on the exposed face of a split embryo (B) and CLSM medial z-planes (C, D). The zone of cell ingression in (B) is colored in purple; the mass of ingressed cells in orange. The ‘belt’ area of the blastocoel wall, colored in green, aligns with the front of the ingressed cells mass. In (C, D) this front is labelled with white arrows; cyan arrows point at individual cells migrating over the blastocoel wall; yellow double-headed arrow indicates the preponderant direction of cell movements within the cell mass. These cells intercalate between each other, aligning as layers with their individual long axes perpendicular to the embryo oral –aboral axis. (E) CLSM of the oral area of an embryo at the mid-gastrula stage. Bottle cells and individual cells of the cell mass are highlighted in turquoise. (F - I): SEM (F, G) and TEM (H, I) of migrating cells. (F) The aboral-most portion of the mass of ingressed cells. (G) The marginal cells of the cell mass move over the blastocoel wall. (H) Sections of intercalating cells flattened perpendicular to the embryo oral –aboral axis. Individual migrating cells are highlighted in orange and violet; a blastocoel wall cell is highlighted in green. (I) Spot-like sites of adhesion between the migrating cells (green arrowheads); yellow arrowhead points to the filopodia-like protrusion. (J, K, L): CLSM (J), SEM (K) and TEM (L) of the ‘belt’ area. The marginal cells of the cell mass (cyan arrowheads) move over the basal surfaces of the blastocoel wall cells (yellow arrows show the direction of this movement). The blastocoel wall cells in the belt area are highlighted in blue in (J) and in green in (K, L); these cells are bent, and their basal ends slant up towards the oral pole. (M) - CLSM of the blastocoel roof; orange arrows point to cells with basally located nuclei. Scale bars: A - D - 50µm; E, J, M - 20µm; F, G, L - 10µm; H - 5µm; I - 1 µm; K - 4µm.

The non-ingressing cells of the blastoderm also show distinct morphologies during mid-gastrula stages. In lateral regions, forming a belt at the level of the underlying front of cell ingression, they become slanted or skewed, with their apicobasal axes bent towards the embryo oral pole (Figs. 4Ce; 7B, J, K, L). We hypothesize that this results from the pulling forces exerted by prospective endoderm cells migrating along the underlying blastocoel roof (see Discussion). Another distinct behavior of blastoderm cells, observed across all of the rounded aboral part of the gastrula, results in its stratification during mid and late gastrula stages. The nuclei of a scattered cell sub-population displace basally as gastrulation proceeds, such that a layer of nuclei forms progressively on the basal side of the blastoderm (Figs. 4Dh; 7M; 8M). It appears that these nuclei belong to a distinct layer of flattened cells lining the blastocoel, but we cannot rule out that thin apical projections connect these cells to the embryo surface. Finally, some mitotic blastoderm cells are observed throughout gastrulation, typically rounding up and adopting apical positions during division (Fig. 8E).

**Figure 8.**
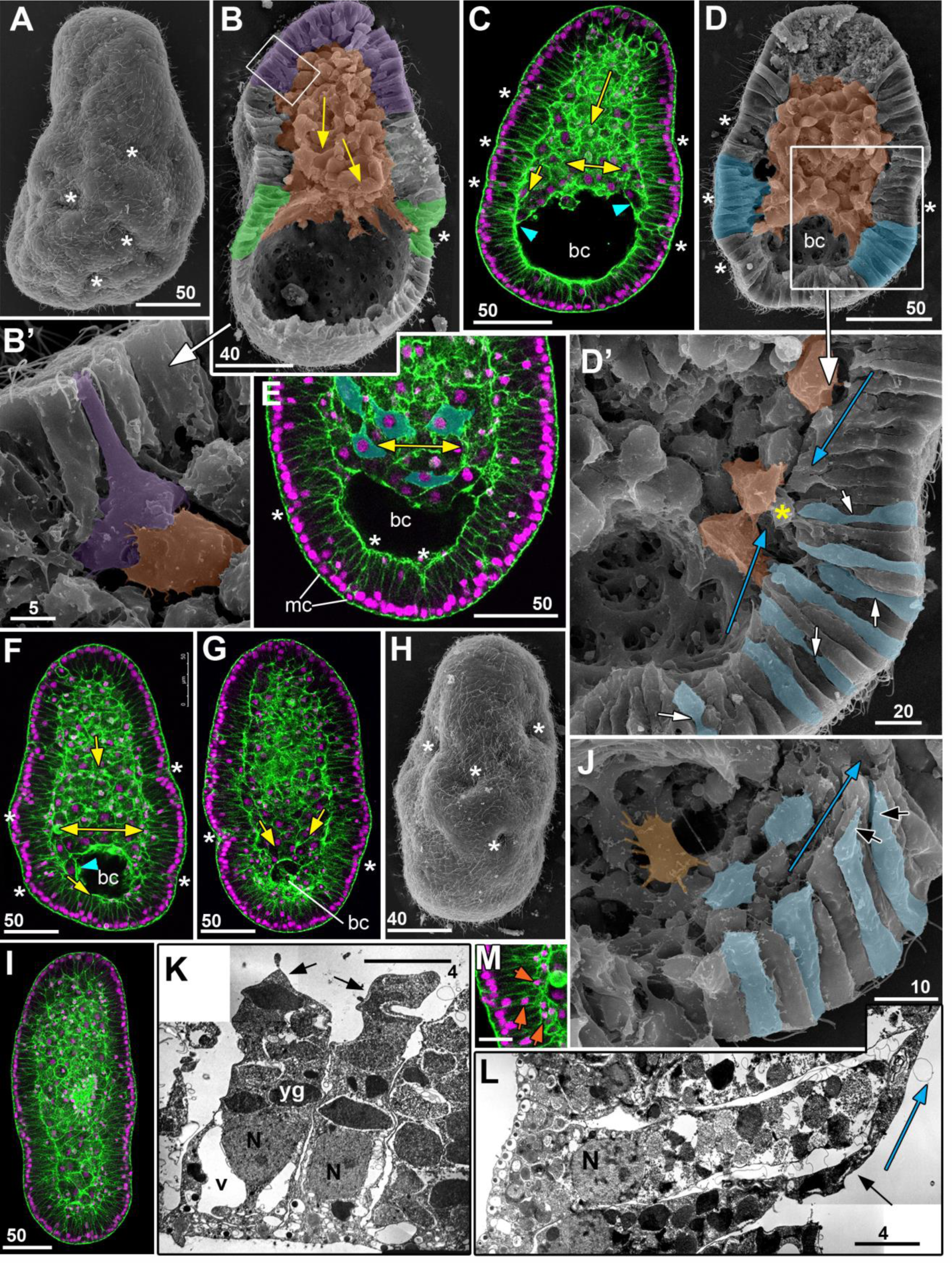
The late gastrula stage and transition to parenchymula stage. (A) SEM of an embryo at the late gastrula stage. Note multiple wrinkles and folds (white asterisks) at the embryo surface, especially the aboral half. (B) SEM image of a split late gastrula stage embryo. Yellow arrows show the direction of cell movements in the mass of ingressed cells, highlighted in orange. A fold-rich region of presumptive ectoderm cells, the belt area, is highlighted in green and the oral pole region of cell ingression in purple. (B’) High magnification of the area framed in (B). Cell ingression is still in progress. A bottle cell is highlighted in violet and an individual ingressed cell in orange. (C - G) Successive stages of blastocoel reduction imaged by CLSM (C, E, F, G) and SEM (D, D’). Note that only aboral front cells of the cell mass align as layers (double-headed arrows on C, E, F), and other cells form streams oriented towards the aboral pole (yellow arrows in C, F, G). Marginal cells of the cell mass move over the blastocoel wall outpacing the front of the cell mass (cyan arrowheads in C, F). (D, D’) SEM of a fold-rich region near the front of the cell mass. Cells of two folds aligned with the front of underlying migrating cells are highlighted in blue. (D’) High magnification of the area framed in (D). Cells of different shapes (side view) are highlighted in blue. Blue arrows show the tilt of the apico-basal axes in cells of a fold. The inner tip of the fold is marked with a yellow asterisk. White arrows point to the cells with expanded basal parts and narrow necks connecting the basal and apical parts. (H) SEM of an embryo at the end of the blastocoel reduction (transition to the parenchymula stage). The shape difference between the oral and aboral halves is less pronounced than in (A). (I) CLSM of an embryo at the parenchymula stage with the blastocoel completely filled with ingressed cells. (J - M) Morphology of the presumptive ectoderm cells at the late gastrula stage. (J) SEM image demonstrating the cell basal surfaces. Black arrows point to elongated basal ends slanted to the tip of a fold. Individual cells lining the blastocoel highlighted in orange arise from the migrating cell mass. (K) TEM of the aboral cells of the same stage embryo as shown in (B); black arrows point to the cell basal ends. (M) TEM of the aboral cells of the same stage as shown in (D, J). Note the difference between the cell apico-basal lengths and cell shapes in (K) and (M). (L) CLSM of a fragment of the presumptive lateral ectoderm. Orange arrows point to the nuclei displaced to the basal area. Scale bars: A, C, E, D, G, H, I - 50µm; B, F - 40µm; B’ - 5µm; E’, L - 20; I - 10µm; K, M - 4µm.

### Late gastrula stage

At the late gastrula stage (Fig. 8; schematized inFig. 4D, E) gross morphological differences between the oral and aboral parts of the embryo progressively diminish (compare Fig. 8A, B with 8G, H). During this period the embryo surface becomes very uneven and covered with wrinkles (Fig. 8A- D, F, G). Around the oral pole, the ratio of bottle cells to epithelial cells gradually decreases as gastrulation progresses, however bottle cells were found even at the late gastrula stage (Fig. 8B, B’), indicating that they continue to be generated over an extended time period in *Clytia*. Mesenchymal-type cells aligned perpendicular to the oral-aboral axis could still be detected in the central core of ingressing cells throughout gastrulation, but become progressively fewer as the embryo narrows and are mainly confined to the front of the advancing cell mass (Figs. 4Dd, Dg, Cg; 8C, E, F). Conversely, flattened cells aligning with and migrating upon the blastocoel wall remain a major feature until gastrulation is complete (Fig. 8C, F, J). In the blastoderm, the abundant wrinkles overly regions or migrating cells (Fig. 8D, D’) and some blastoderm cells slant into the wrinkles from both sides (Figs. 4Ei; 8D’, J, L). The end of cell ingression, at about 24hpf, marks the parenchymula stage, from which point the embryonic surface morphology becomes smooth again (Figs. 8I and 9A, B).

### Planula development from the parenchymula; endodermal cell epithelialization

During the 24 hours following the completion of cell ingression, the main event of planula morphogenesis is the reorganization of ingressed cells to form an endodermal epithelial layer: ingressed cells close to the blastocoel wall reorganize to form a new internal epithelium of columnar, polariz ed cells (Figs. 4F, G; 9A-D, 9G, G’, H, H’). This has two important consequences for tissue organization. One is the formation of a gastrocoel cavity. This process is accompanied by death of cells in the endodermal region of the planula, with fragmented nuclei accumulating in the central core where the cavity develops (Fig. 9A, C, D, G’, H, H’). We performed TUNEL staining to detect nuclei in which DNA is beginning to fragment ahead of cell death. This revealed that dying cells are dispersed amongst the endodermal layer as the gastrocoel forms, their numbers increasing between 1 and 2 days post fertilization (Fig. 9N, N’, Supplementary Fig. S4). A likely interpretation of this distribution is that cells that fail to incorporate correctly into the endodermal epithelium undergo cell death and degrade in the central cavity to be digested by the endodermal layer. The role of cell death in contributing to gastrocoel formation remains to be tested . In most planulae examined, TUNEL stained and fragmented nuclei were relatively scarce, but occasional specimens showed a much greater accumulation of dying cells in the endodermal region (Supplementary Fig. S4D), likely derived from cells that fail to develop correctly during the cleavage or blastula stages and detach into the blastocoel.

**Figure 9.**
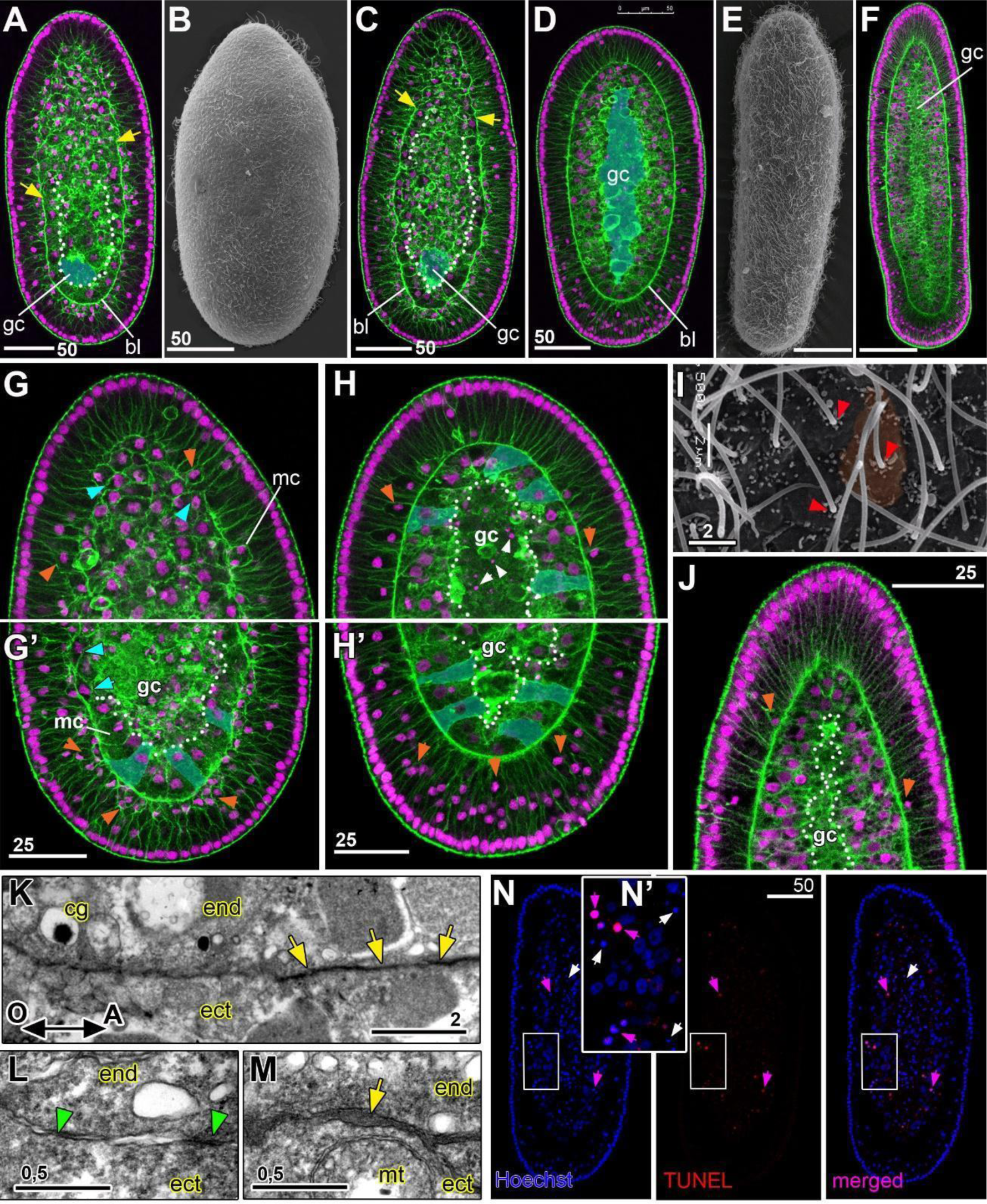
Formation of the planula. Epithelization of the endoderm and cell differentiation. Developing planulae: 1 (A-C, G, G’), 2 (D, H, H’) and 3 (E, F, I, J) days post fertilization with oral poles at the top in all panels. (A, C, G, G’) CLSM of embryos at the beginning of the gastrocoel formation. A small gastrocoel (gc) appears first near the aboral pole (highlighted). The basal lamina (bl) and an epithelial endoderm start to form in the aboral half. Yellow arrows indicate the oral limit of the basal lamina visible on the CLSM section in (A, C); white dots outline the forming enodermal epithelium. (C) In this embryo the basal lamina has extended further towards the oral pole than in (A). (G, G’) High magnification of the oral (G) and aboral (G’) areas of the embryo shown in (C). Cyan arrowheads point to endodermal cells with a spindle shape in cross-section characteristic of migrating cells. Orange arrowheads point to the ectodermal cells with the basally located nuclei. Note the basally and apically localized mitotic cells (mc) in (G) and an endodermal mitotic cell in (G’). Individual epithelial cells in the endoderm are highlighted in blue in (G’). (D) Early planula (2 days post fertilization). Gastrocoel formation and endoderm epithelization are complete. (H, H’) High magnification of the oral (H) and aboral (H’) areas of the embryo shown in (D). White arrowheads indicate nuclear fragments in the central core region. Individual epithelial cells in the endoderm are highlighted. (E, F) Mature planulae (3 days following fertilization) with an elongated body, pointed oral (posterior) pole and rounded aboral (anterior) pole. (I) Apical surfaces of the planula ectoderm cells. One of these cells is highlighted in orange. Red arrowheads point to the cilia. All the cilia are positioned towards the aboral (anterior) side of the cells. (J) High magnification of the planula oral (= posterior) pole. (K - M) TEM of the border between ectoderm (ect) and endoderm (end) in 1 dpf embryo (parenchymula). (K) Lateral area of an embryo demonstrating the aboral - oral gradient of basal lamina formation; yellow arrows indicate a well formed part of the basal lamina; cg - cortical granule in the endoderm cell. (L) Higher magnification of an oral area where the basal lamina is not yet formed; green arrowheads show the sites of adhesion between the ecto- and endoderm cells. (M) Higher magnification of the area with the developed basal lamina; mt - mitochondria. (N) Visualization of TUNEL stained DNA (magenta) in 2dpf planula; magenta arrowheads illustrate TUNEL-positive fragmented nuclei amongst cells of the endodermal layer; White arrows - TUNEL-negative nuclear fragments in the central core region. (N’) Higher magnification of the framed area. Further examples in Supplementary Fig. 3. Scale bars: A, F, N - 50µm; G, G’, H, H’, J - 25µm; I, K - 2µm; L, M -0,5µm.

A second consequence of endodermal epithelization is the appearance of a distinct basal lamina separating the ectodermal and endodermal epithelia (Fig. 9A, C, D, G’, H, H’, K - M). This basal lamina stains strongly with phalloidin because of actin-rich contractile projections that develop on the basal surface of the differentiating epitheliomuscular cells that predominate in both layers. By TEM, basal lamina appears as a thick layer of ECM separating the ectodermal and endodermal cells (Fig. 9K, compare 9L and 9M). As for gastrocoel formation, endoderm epithelization and basal lamina formation start from the aboral pole and proceed orally (compare Figs. 9G and 9G’; 9G and 9H).

Planula larva formation is completed by approximately 48hpf (Fig. 9D). During the next 24 hours, morphological differences between the oral and aboral ends of the planula become more pronounced (Fig. 9E, F, J). Cilia on each ectodermal cell of the planula are clearly positioned towards the cell oral side (Fig. 9K) as described previously (Momose et al., 2012). In the course of planula formation, the blastoderm transforms into a more complex ectodermal epithelium with richer variety of cell types (nematocytes, neural cells and secretory gland cells) (Bodo and Bouillon, 1968). This process is completed between two and three days after fertilization.

### Gene expression changes accompanying gastrulation

To help understand how the cell morphological changes observed during gastrulation relate to regionalized expression of regulatory genes we performed in situ hybridization on embryos fixed at successive stages (Fig. 10A). Two *Clytia* brachyury genes, *CheBra1* and *CheBra2,* are expressed downstream of the oral Wnt3 signaling that defines the oral /endodermal domain at the early gastrula stage (Momose & Houliston, 2007; Momose et al., 2008; Lapébie et al., 2014). We detected *CheBra1* and *CheBra2* expression in overlapping cell populations within the oral domain of the blastoderm before and during gastrulation, and finally of the planula ectoderm, but not in ingressed cells (Fig. 10A, B). These two *Clytia* brachyury paralogues appear to have partly redundant functions in gastrulation (Lapébie et al., 2014), but may have more distinct roles in adult stages as shown for their orthologues HyBra1(endodermal) and HyBra2 (ectodermal) in the *Hydra* polyp oral pole region (Bielen et al., 2007). The *Clytia* orthologue of Snail, a transcriptional repressor implicated in EMT as well as cohesive cell migration across many species and developmental contexts (Barrallo-Gimeno & Nieto, 2005), showed a dynamic expression pattern. It was detectable in cells across the entire blastoderm at the late blastula stage, before becoming concentrated more orally at the onset of gastrulation. During these stages, expression of *CheSnail, CheBra1* and *CheBra2* all showed a small local reduction at the oral pole (Fig. 10A, B). In mid-gastrula stage embryos, detectable *Snail* expression became confined to ingressing and internalised cells, notably ones at the front of the migrating cell mass, and by planula stages *CheSnail* expression was only weakly detectable (Fig. 10A).

**Figure 10.**
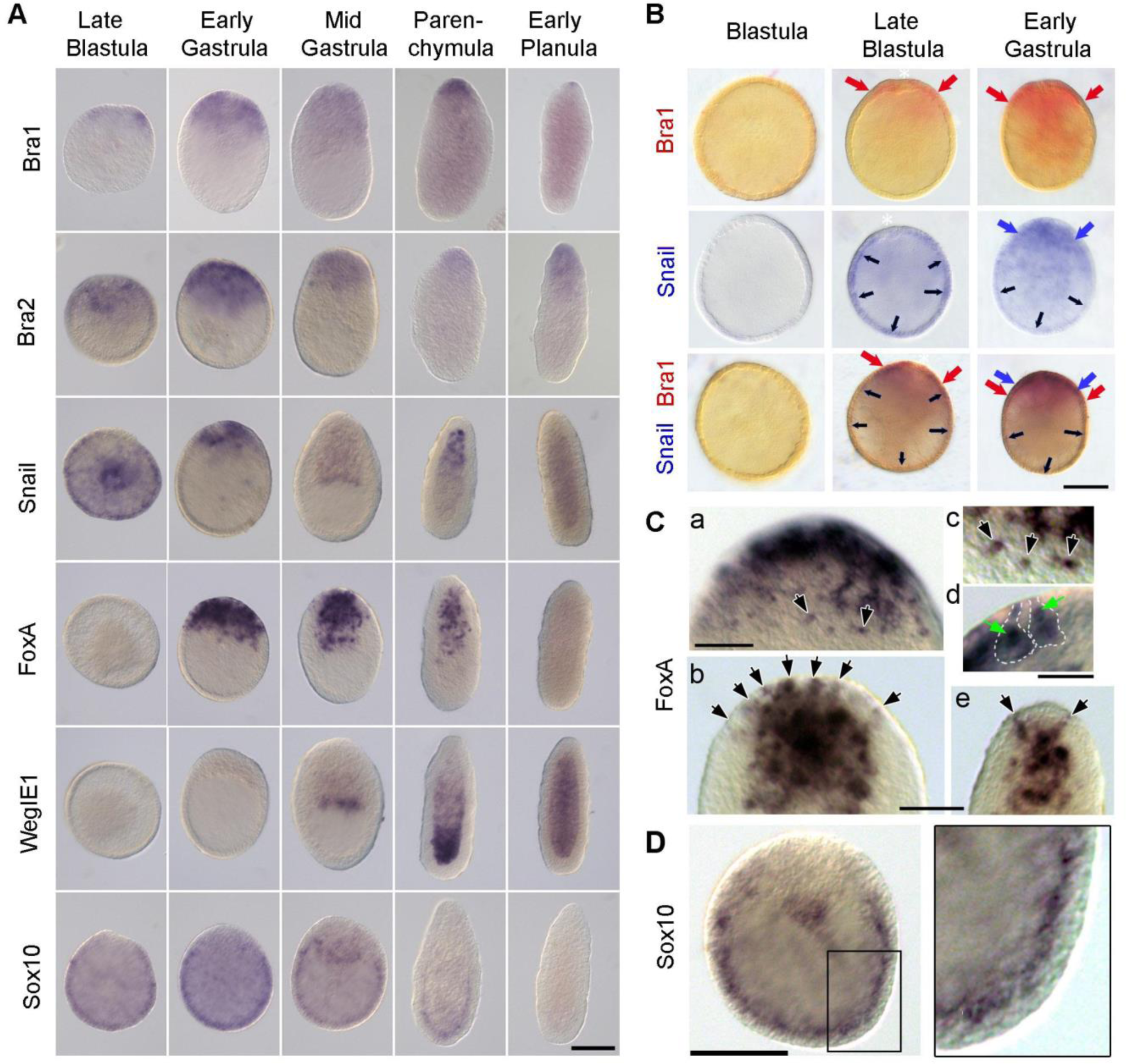
Expression of regulatory and territory marker genes during gastrulation. (A) In situ hybridization detection of *CheBra1*, *CheBra2*, *CheSnail*, *CheFoxA*, *CheWegIE2*, and *CheSox10* mRNAs at the late blastula, early gastrula, mid-gastrula, parenchymula (1 day pf) and planula (2 day pf) stages, as indicated by labels on the left and top of the panel. (B) Double in situ hybridization detection of *CheBra1* (red), and *CheSnail* (blue) expression at late blastula and early blastula stages. Red and blue arrows indicate the aboral limit of *Bra1* and *Snail* expression, respectively. At the blastula stage *Snail* expression is mainly uniform, but ‘holes’ of low expression for each gene coincide at the oral pole (white asterisks). (C) Detection of *FoxA* on early gastrula (a, c, d), mid-gastrula (b) and parenchymula (e) stage embryos. Surface view (a, c) allows detection of the contracted apices of *FoxA* positive bottle cells (black arrows). Optical sections (b, d, e) show the individual bottle cells (black arrows). Two bottle cells are outlined with white dotted lines on (d); their nuclei are shown with the green arrows. (D) Enlargement of *Sox10* mRNA detection through an early gastrula stage embryo, emphasising the position of the stained cells on the basal side of the inner lateral and aboral blastoderm Optical sections are through the embryo center on the left, and at the level of the blastoderm basal layer on the right. Oral pole is up in all panels. Scale bars: A, B, D - 100µm; Ca, Cb, Cd - 50µm, Cc, Cd - 25µm.

Expression of the *Clytia FoxA* transcription factor (Lapébie et al., 2014), whose orthologues are associated with the development of digestive territories in many animal species (Steinmetz et al, 2017), was detected from the early gastrula stage. They initially concentrated in the oral blastoderm and showed bottle-cell like morphology with constricted apical domains (Fig. 10A, Ca - Cd). The ingressed cells retain the *CheFoxA* expression (Fig. 10A, C). The number of *CheFoxA* expressing cells in the oral blastoderm and in the developing endodermal mass diminished progressively as gastrulation proceeded (Fig. 10A, Ce). At the parenchymula stage, FoxA positive cells were detected only in the oral region of the endodermal mass. By the planula stage *FoxA* expression was no longer detectable. The *Clytia*-specific endoderm marker gene *WegIE2* (Lapébie et al., 2014) became detectable in more aboral regions of the internalized cell mass, consistent with a switch in expression between these two genes in the course of endoderm cells differentiation.

Finally, expression of the Sox10, a B-type Sox transcription factor expressed in neural cells of the adult *Clytia* medusa (Jager et al., 2011), was detected throughout the blastoderm except in the oral pole ingressing cells (Fig. 10A, D). Expression persisted in the basal region of lateral and aboral ectoderm cells until the end of gastrulation (Fig.10D). Residual expression could be detected in a basal domain of the aboral pole ectoderm at the parenchymula stage. This Sox10 expression pattern coincides with the accumulation of basal nuclei in the lateral blastoderm during nuclear stratification described in Figs. 7M, 8M, 9G’, suggesting that cells with a neural fate are generated in this position. Correspondingly, Sox10 expression in later stage planula could detected in neural cells concentrated at the poles of the embryo as reported previously (Jager et al; 2011), but required much longer in situ hybridization development times than for gastrula stage embryos, indicating that Sox10 mRNA levels in decreases during neural cell differentiation.

## Discussion

We have reconstructed the changes in cell shape, cellular behavior and underlying ultrastructure associated with successive stages of gastrulation in *Clytia hemisphaerica,* as summarized in Figure 4. These transform a simple blastula comprising an irregular epithelial ball into an elongated, polarized, planula larva with two structured epithelia, endodermal and ectodermal. Gastrulation proceeds by unipolar cell ingression: individual cells ingress asynchronously into the blastocoel from within an oral domain throughout the gastrulation period (Fig. 4B - D; see also Supplementary Movie M2). Ingressed cells migrate towards the aboral pole. As the blastocoel partially fills, the whole embryo elongates and adopts a pear shaped morphology, with characteristic constriction of the developing ectodermal layer in a belt region overlying the front of cell migration (Fig. 4C, D). The mid- and late gastrula stage embryos demonstrate pronounced wrinkling of the lateral blastoderm associated with concerted bending of the basal domains of the epithelized cells (Fig. 4D, E). Finally, as the parenchymula adopts its smoother, pointed torpedo shape, the inner mass of cells reorganize into an epithelial layer in an aboral-oral progression, and an extracellular mesoglea is elaborated to separate the basal surfaces of the two epithelia (Bodo and Bouillon, 1968; Fig. 4F, G). The elaboration of the epithelial layers through cell differentiation also progresses from aboral to oral in hydrozoan larvae (Gröger & Schmid, 2001): myofibrillar foot processes that extend from differentiating epithelial cells, intermix with neurites projecting from neurosensory and ganglionic cells (Lapébie et al., 2014; Thomas et al., 1987).

The morphology of a developing embryo and of the cells within it are intimately linked. The particular mechanical environment of the embryo influences how individual cells behave, while conversely cell behavior influences the mechanics of tissues and of the whole embryo (LeGoff & Lecuit, 2015). Indeed, mechanical forces can be considered among the primary coordinators of morphogenetic processes, with cell behavior being guided by forces transmitted over long range (Heisenberg & Bellaïche, 2013). Below we attempt to interpret embryo morphology during *Clytia* planula formation in terms of forces and tensions generated by collective cellular movements and shape changes of distinct cell sub-populations

### Blastoderm epithelization

The initial morphogenetic processes of *Clytia* development are blastula formation through successive cleavage divisions, and epithelialization of the blastula (blastoderm) cells. Epithelia are a distinctive feature of Metazoa, consisting of sheets of clearly polarized and aligned cells with elaborate cell–cell junctions (Knust & Bossinger, 2002), and are the first tissues that form during embryogenesis (Tyler, 2003). Adult epithelia are generally supported by an extracellular matrix on the basal side called the basement membrane or basal lamina. In adult cnidarians, the characteristic outer and inner epithelial body layers are separated by back-to-back basal laminae, often continuous with a central fibrous matrix of ‘mesoglea’. Cnidarian basal laminae (= basement membranes) resemble those described in bilaterian species on both morphological and molecular levels (Barzansky et al., 1975; Pedersen, 1991; Tucker & Adams, 2014; Buzgariu et al., 2015).

Blastoderm epithelization in *Clytia* generates a cell layer possessing a ‘minimal set’ of epithelial characteristics, with a distinct basal lamina developing only at the planula larva stage as in other cnidarian embryos (Fritzenwanker et al., 2007; Kraus et al., 2014). Similarly, embryonic epithelia in other animals may lack adult epithelial features. For instance, blastoderm cells in the nematode *C. elegans* possess apico-basal polarity but have no discernable basal lamina.

In *Clytia*, morphological signs of apico - basal polarity appear at the mid-blastula stage (7-8 hpf) and become more pronounced at the late blastula stage (9-10hpf). At the same stage (mid-late blastula), blastoderm cells start to acquire planar polarity, the second type of polarity characteristic for epithelial tissues (Momose et al., 2012). Thus, from the late blastula stage, *Clytia* embryo has a functional and mechanically coherent epithelium. Such an embryo is capable of ‘global’ regulation of morphogenetic processes, i.e. cell behavior and cell shape changes are coordinated across the whole embryo by the coupling of molecular and mechanical signals.

### Cell ingression

Cell heterogeneity first arises across the whole embryo at the blastula stage (Fig. 4A). Elongated cells with a bulbous basal region arise, interspersed with shorter cells tapered to a basal point (“squat cells”). The behavior of “basally expanded” cells differs between regions. In the oral domain, they elongate further, transform into bottle cells and eventually ingress into the blastocoel (Fig. 4H). In aboral/lateral domains, these cells do not show increased elongation along the apicobasal axis, and so do not become true bottle cells.

Cell ingression during *Clytia* gastrulation is an example of Epithelial - Mesenchymal Transition (EMT). During EMT, epithelial cells acquire features of mesenchymal phenotypes and behaviour and downregulate/lose epithelial features, the details varying from system to system (Shook, Keller, 2003; Nieto et al., 2016; Campbell & Casanova, 2016; Yang et al., 2020). EMT of cells in the primary embryonic epithelium (blastoderm) has been termed ‘primary EMT’ (Shook, Keller, 2003; Lim, Thiery, 2012; Das et al., 2018). The starting point of primary EMT varies from well-organized epithelia as in the sea urchin blastula, to cell layers possessing only a subset of epithelial features (e.g. apico-basal polarity and intercellular contacts) as in the nematode or frog embryo (Shook, Keller, 2003). The end point also varies, along a continuum of cell states between epithelial and fully mesenchymal phenotypes. Furthermore, the behavior of cells undergoing EMT may vary from individual migration to collective migration as a cohort, in which cells maintain adhesion contacts (Campbell & Casanova, 2016). Despite the variability of initial and final stages of primary EMT, typical stages of cell morphology changes can be distinguished across many species (Shook, Keller, 2003). Very often (but not always) primary EMT proceeds through formation of bottle cells, preceded by breaking down of the basal lamina if present (e.g. in mouse and chick gastrulation). Concurrently, cell apices start to constrict, minimizing the area of cell apical adhesion and preventing the formation of a big hole in the epithelium (Shook, Keller, 2003). A forming bottle cell elongates the apico-basal axis forming the neck connecting the apical and basal parts of the cell; cytoplasm and organelles (including the nucleus) displace to the widening basal part of the cell. The final features of primary EMT are detachment of bottle cell apex from neighboring cells; ingression of the cell from the epithelium (by shedding or active migration); and quick sealing of the wound left in the epithelium by neighboring cells, which form multiple protrusions covering the wound and establishing new intercelluar junctions. These various EMT features do not show a conserved sequence nor necessarily proceed to the final stage (Shook, Keller, 2003; Campbell, Casanova, 2016; Nieto et al., 2016).

There are obvious gaps in our knowledge on primary EMT in cnidarians. While formation of bottle cells was described in many cnidarian species in XIX century (Metchnikoff, 1886), and diverse gastrulation modes have since been described (Kraus and Markov, 2017), ultrastructural and molecular data on EMT have been obtained for only two species: the anthozoan *Nematostella* (for review see Technau, 2020) and the hydrozoan *Clytia* (this study). In *Nematostella* and *Clytia*, primary EMT proceeds through the classic stages of bottle cell formation described above (Shook, Keller, 2003) and illustrated for *Clytia* in Figs. 6B, 4H). Bottle cells in both species express *Snail*, the key transcription regulator of EMT. However, in *Nematostella*, EMT arrests prior to the final stage and cells do not ingress (Magie et al., 2007). The presumptive endodermal cells acquire a migratory phenotype and behavior while still connected by adherence junctions. Their apical cell surfaces are conserved as the apical perimeter constricts and cilia are retained (Kraus, Technau, 2006; Technau, 2020). In contrast, presumptive endoderm in *Clytia* ingress individually into the blastocoel detach and migrate in the manner of mesenchymal cells. They show an important feature of primary EMT by retaining intercellular junctions almost until the moment of their release from the epithelium, while the area of these contacts gradually decreases (Katow & Solursh, 1980; Nakaya & Sheng, 2009). The mechanisms responsible for junction loss remain to be established.

No remnants of intercellular junctions were observed on the membrane of the trailing edges of ingressed cells, nor did we observe endocytotic vacuoles containing fragments of junctions. *Clytia* bottle cells also retained their cilia until the final stages of EMT. The cilium sinks into a deep pocket of the cell apical membrane (Fig. 6G), implying that it becomes internalized within the cell rather than shed from the cell surface. Loss of cilia during primary EMT has also been observed in other species. The underlying mechanisms have yet to be investigated, although in sea urchin embryos, cilia fragments were found inside the vacuoles of ingressed cells (discussed in Katow & Solursh, 1980).

Upon ingression, *Clytia* cells maintain a mesenchymal morphology and migrate towards the aboral pole in a highly coordinated manner (Figs 4 B - D; 7E - J; 11A - C). They are connected by spot-like sites of adhesion (Fig. 7I) which resemble the cell-cell contacts characteristic for vertebrate mesoderm cells migrating as a cohort (Campbell & Casanova, 2016). The coordinated behavior of the ingressed cells likely accounts for the global pear-shaped morphology of mid- gastrula stage embryos and torpedo-shaped morphology of parenchymula stage embryos. Intercalation and coordinated aboral migration of these cells reduces the diameter of the oral region first, and, finally, of the whole embryo. Consistently, disruption of cell intercalation orthogonal to the oral-aboral axis by morpholino-knockdown of PCP proteins (Momose et al., 2012) or as modelled in silico (van der Sande et al., 2020), prevents the narrowing of the embryo during gastrulation.

These comparisons together illustrate how the features of primary EMT, which provide the integrity of embryonic epithelia as well as the migration and cohesion of ingressing cell, are shared between Cnidaria and other animal species and likely can be traced to the roots of Metazoa.

### Cell morphology diversification and gene expression

Computational models indicate that the emergence of two distinct cell forms from an initially uniform epithelial population during is driven by apical constriction occurring in all blastoderm cells during compaction in *Clytia* (van der Sande et al., 2020), and in cells of the oral domain (presumptive endoderm) at the onset of gastrulation in *Nematostella* (Tamulonis et al., 2011). “Basally expanded” cells located in the oral domain likely provide the basis for bottle cell formation in both species.

During gastrulation some oral pole cells remain at the oral pole to sculpt its characteristic point, while others ingress into the blastocoel to form the endoderm. During this morphological segregation, the former population maintains expression of the two *Clytia brachyury* genes and the latter *CheSnail*. This fits with a general theme in animal gastrulation of Snail acting in ingressing and migrating cells, and Brachyury in epithelial cells around at the gastrulation site. Brachyury proteins in animals are thought to have retained roles in cell morphogenesis based on regulating actin-based cell motility, shape and behavior from a unicellular ancestral transcriptional activator (Sebe-Pedros & Ruiz-Trillo, 2017). They have also acquired functions in determining mesoderm or endoderm tissue fates, and in defining organizing centers through the cross-regulation with Wnt-beta catenin and other signaling pathways, which have complicated interpretation of their exact roles in morphogenesis from knockdown studies (Technau 2001; Hejnol & Martin-Duran 2015; Servetnick, et al, 2017). Snail transcriptional repressors are widely involved in regulation of cell adhesion and migration during gastrulation, including EMT of presumptive mesoderm cells from the chick epiblast and collective cell migration of fish involuting mesendoderm cells (Barrallo-Gimeno & Nieto, 2005; Lim & Thiery, 2012). Thus, during EMT–based germ layer segregation processes such as primitive streak formation and tailbud paraxial mesoderm formation in amniotes, as in *Clytia* gastrulation, Brachyurys co-express with Snails or other EMT/ cell-motility regulators (eg Sox4 and Bves) prior to cell ingression, but not in ingressed cells (Kispert et al., 1995; Lolas et al., 2014; Goto et al., 2017). For species in which gastrulation involves invagination or involution of epithelial layers, including anthozoan cnidarians, amphibians, ambulacraria as well as diverse protostome species, brachyury family genes are expressed in cells associated with the blastopore lip (Scholz & Technau, 2003; Gross and McClay, 2001; Evren et al., 2014; Sebé-Pedrós & Ruiz-Trillo 2017). Experimental interference with them disturbs blastopore morphogenesis in some but not all cases (Gross and McClay, 2001; Evren et al, 2014; Yasuoka et al 2016; Servetnick et al., 2017). Brachyury expression at the blastopore lip is itself reinforced mechanically by local cell deformations at the epithelial bend in a beta-catenin signaling context (Brunet et al., 2013; Pukhlyakova et al., 2018). Similarly, *HyBra1* and *HyBra2* expression during budding in *Hydra* polyps is localized to bends in epithelium of future oral tissues, and as in the gastrulation context occurs within a domain of beta catenin signaling. Bracyurys are similarly expressed at oral “lip” positions around mouth openings in *Hydra* and other animals (Bielen et al. 2007; Gross and McClay, 2001; Hejnol & Martin-Duran 2015). A computational model for *Nematostella* gastrulation predicts that strain at the blastopore lip is promoted by reduction in cell adhesion in the adjacent domain of presumptive endoderm (Tamulonis et al., 2011), which is marked by *Snail* expression (Fritzenwanker et al., 2004; Hayward et al., 2015). Snail likely contributes to the observed “partial EMT” and progressive transformation of these oral-most cells into endoderm by repression of Par and cadherin gene expression (Fritzenwanker et al., 2004; Botman, et al., 2014; Salinas-Saavedra et al., 2018; Pukhlyakova et al., 2019). We can propose an equivalent mechanism in *Clytia* based on the predictions of a computational model in which a *Nematostella*-like gastrulation mode was converted into an ingression mode by increasing the extent of adhesion loss in the oral domain while maintaining apical cell constriction across all cells (van der Sande et al., 2020). We propose that as weakened adhesion driven by *Clytia Snail* expression causes ingression of bottle cells, resultant strain on the interspersed ectoderm ‘squat’ cells maintains Brachyury gene expression in the oral ectoderm. The *Snail* expressing cells undergo complete EMT, before migrating together aborally as a cohesive unit. This hypotheses could be explored in future studies by cellular resolution detection of *Bra1, Bra2* and *Snail* combined with mechanical manipulations, as well as studies of the associated gene regulatory networks and signaling pathway (Lapébie et al, 2014; Abdol et al., 2017).

FoxA (=Forkhead) expression profile comparisons between species are suggestive of involvement in fate determination rather than mainly in morphogenetic movements. *NvFoxA* is expressed early during blastopore formation in the same domain as *NvBra* (Fritzenwanker et al., 2004; Magie et al., 2007), later giving rise to the pharyngeal ectoderm. Cells of his domain will develop exocrine digestive functions, linking it evolutionarily to endoderm domains in bilaterian species (Steinmetz, 2019). *Amfkh* is only transiently expressed in the developing blastopore, this being spatially and temporally separated from its later expression in the pharynx (Hayward et al., 2015). In *Clytia*, FoxA is transiently expressed in bottle cells (Fig. 10C), but the fate of these cells is not yet known. It will be interesting to determine whether these FoxA expressing cells give rise to any particular endodermal cell type.

The “basally expanded” cells in lateral/aboral regions (presumptive ectoderm) of the *Clytia* early gastrula embryo do not transform into bottle cells, but rather contribute to a distinct basal layer of the blastoderm via a process of stratification /pseudostratification. This idea remains to be demonstrated experimentally, but is supported by the transient expression in basal regions of the blastoderm of the transcription factor Snail, and subsequent expression of the putative neurogenic SoxB family transcription factor Sox10 (Fig. 10D; Jager et al., 2011). Basal cells of the lateral/aboral blastoderm may thus provide precursors of particular neuronal cell types (“ganglion cells”) that can develop from aboral gastrula fragments (Thomas et ale., 1987). Generation of a basal blastoderm layer has also been observed in the hydrozoan Gonothyraea loveni, in this case via oriented cell divisions (Burmistrova et al., 2018).

### Forces and tensions sculpting embryo morphology during gastrulation

Mid and late stage gastrula embryo provide a complex landscape for inferring tissue forces and tensions from embryo and cell morphologies, as schematized in Fig. 11. As the ingressed cells fill the blastocoel, they form three populations: a “shell” of tightly-associated flattened cells crawling along the blastocoel wall (Figs 7G, J, K; 11B, C), flattened intercalating cells in the “core” oriented perpendicular to the oral-aboral axis (Figs 7E, F, H; 11B, C), and a less ordered core population closer to the oral pole (Fig. 11B, C). In concert the embryo shape changes markedly - the oral area progressively narrows and becomes pointed (Figs 7A; 8A, H; 9B, E; Fig. 11 Ba, Ca), while the locally constricted “belt” zone of the blastoderm accompanies the front of underlying cells migrating towards the aboral pole (Figs 7B, C; 8B; 11Ba, Bb). We propose that embryo morphology is dictated primarily by the shell cells and the intercalating core cells. The migrating shell cells are predicted to generate the blastodermal folds beneath and behind them, as well as tension lines in front of them that in turn guide migration. These cells associate closely with each other (see spots of adhesion in Fig. 7I) and appear to migrate aboralwards as a cohesive structure (dark blue arrows in Fig. 11Bd, Ce). The coherent movement of the cells at the aboral front of migration resembles that of ‘leader’ cells during wound healing (du Roure et al., 2005; Poujade et al., 2007). Forces generated by the actin cytoskeleton of migrating cells generate wrinkles in the substrate (Fournier et al., 2010), in this case on the blastocoel wall (Fig.11 Bd, D, E). Compression of the *Clytia* blastocoel wall beneath and behind the migrating cells into a mass of microscopic folds, oriented perpendicular to the direction of cell migration (Fig. 11 C, F), could result from amplification of wrinkles as accumulating deformations create additional mechanical stresses (Lange & Fabry, 2013). In contrast, the blastocoel wall ahead of the collectively migrating cells becomes stretched into radiating folds oriented along the lines of tension (Fig.11 D, E, Cb, Ce), as has been demonstrated using artificial deformable substrates (Harris et al., 1980; Pelham & Wang, 1999). These lines of tension are good candidates to orient the migration of the shell cells along the oral-aboral axis (Lange & Fabry, 2013; Tambe et al., 2011).

**Figure 11.**
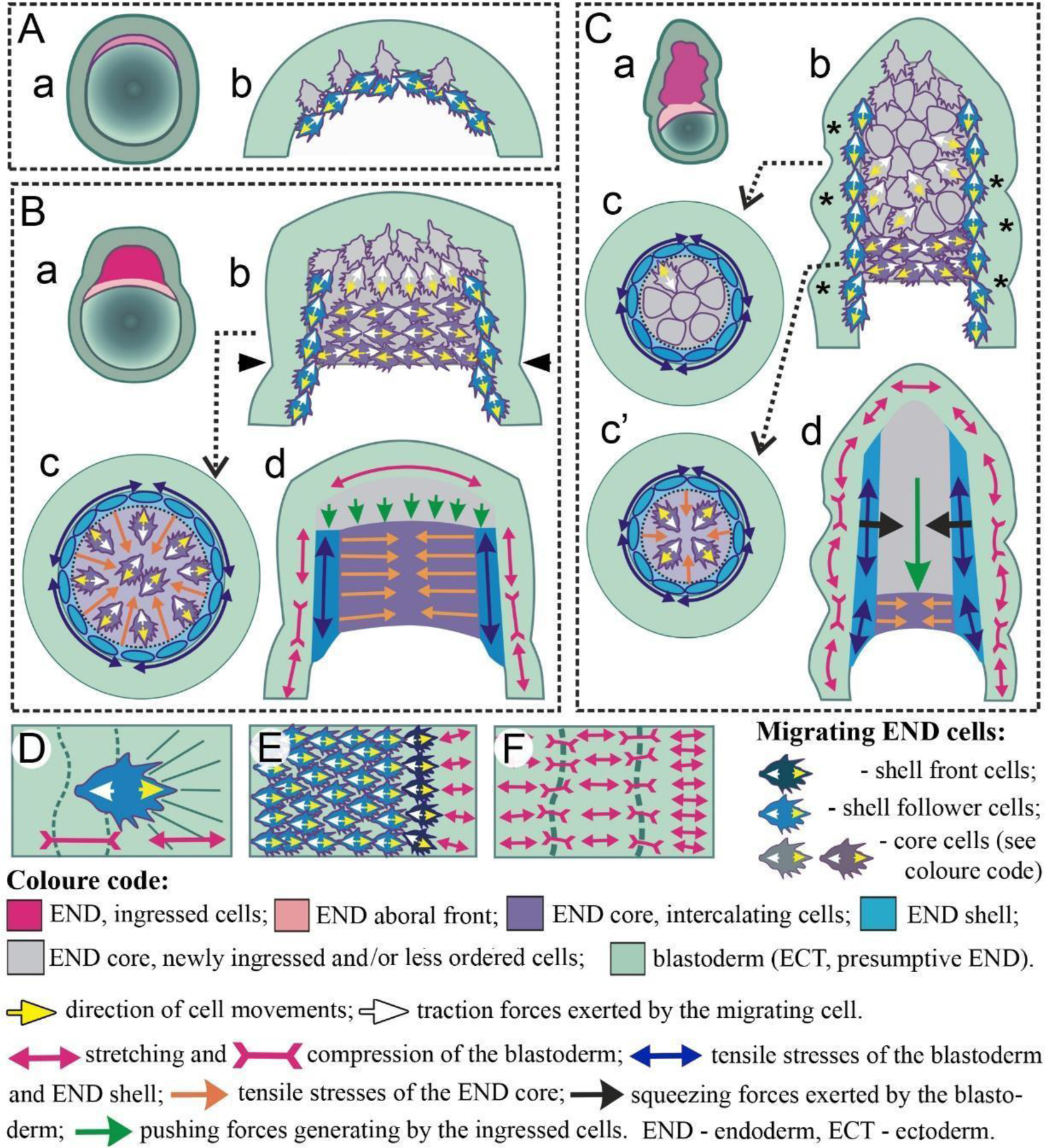
Interpretation of embryo and cell morphology during *Clytia* gastrulation in terms of forces and tensions. The composite panels illustrate for successive stages of gastrulation the spatial distribution of cell morphologies, cell movements and predicted forces and tensions driving morphogenesis. A - early gastrula, B - mid-gastrula, C - late gastrula. Individual cartoons show embryo midsections with oral at the top (a), zooms on the oral end (b, d), cross sections at the levels indicated (c and c’), and predicted forces and tensions (c, c’, d). Dashed lines in c and c’ cartoons indicate the frontier between the ‘shell’ of crawling cells and the inner ‘core’; arrowheads in B indicate the ‘belt’ area of the blastoderm. Macroscopic folds of the blastoderm in C are marked with the asterisks. Panels D-F illustrate examples of the relationship between individual cell behavior and force predictions. (D) An individual ingressed cell of the ‘shell’ migrating over the blastoderm basal surface generates traction forces oriented opposite to cell migration and acting on the substrate. The blastoderm beneath the cell on its trailing side is compressed into micro-folds (dashed lines) perpendicular to the direction of motion, and ahead of migration is stretched into the radial wrinkles (solid lines). (E) Multiple cells migrating over the blastocoel wall at the mid/late gastrula stage similarly induce stretching of the area above the front of migration. Cells of the aboral front of migration are colored in dark blue. (F) Blastocoel wall surface; migrating cells were removed to show alternating areas of stretching and compression. Macroscopic folds (dashed lines) correspond to the areas of compressiOn. The colored arrows represent predicted cell forces and tensions as indicated in the key.

The second cell population of key importance to gastrula morphogenesis comprises flattened mesenchymal cells stacked along the oral-aboral axis towards the front of the advancing core (dark violet cells in Figs. 11 B, C; 4Cd, Dd). Their organization and morphology is highly reminiscent of involuting mesoderm cells that intercalate towards the midline during chordate gastrulation (Wallingford et al. 2002). In both cases, narrowing of the internal internalizing cell mass and the associated embryo elongation, is disrupted by knockdown of the PCP proteins (Momose et al., 2012). Our 2D model of *Clytia* gastrulation (van der Sande et al, 2020) indicates that lateral crawling interactions between the ingressed cells make an important contribution to embryo elongation. PCP-depended lateral intercalation between cells of the blastoderm also likely contributes (Byrum, 2001; Momose et al., 2012). Another consequence of intercalation between ingressing endoderm cells, also suggested by Byrum (2001), is the narrowing of the oral region during gastrulation. Forces generated in the front portion of the ingressing mass pull the blastocoel wall towards the center leading to formation of the constricted “belt” (Fig.11 B).

A third, more disordered, cell population arises in oral regions of the core during gastrulation as further cells ingress (grey areas in Fig. 11 Bb, Bd, Cb, Cd). We propose that most of these cells initially join the core rather than the shell because the layers of aligned core cells are under tension and thus attract newly ingressed migrating cells (Lo et al., 2000). By the end of gastrulation, cell alignment in the endodermal core is less pronounced, as the core becomes narrower perpendicular to the oral-aboral axis and relatively higher number of cells become peripheral cells, which align with the blastocoel wall (Figs. 8C, G; 11Cb, Cd). Ingressing cells will rather join the stiffened shell that is under the strong tension at this stage.

Thus, the much reduced diameter of the parenchymula reflects both loss of cells from the blastoderm and lateral intercalation of the core cells. Diameter reduction in turn leads to reduction of tension inside the core (Fig.11 Cd). At the same time, diameter reduction generates squeezing forces perpendicular to the OA axis and contributes to the aboralward displacement of ingressed cells (Fig.11 Cd). The polarized morphology of the canonical hydrozoan larva, including its pronounced elongation and tapered posterior end emerges towards the end of gastrulation (Fig.11 C).

Future studies will be required to test these predictions about forces tensions and cell behavior. One approach would be to target the molecular and cellular mechanisms of both individual cell shape changes and overall physical properties. For instance, adhesion molecules and ECM components contributing to the cohesive migratory behaviors of the cells could be tested by downregulation of candidate cell surface molecules expressed in these cells, or signaling molecules expressed in the oral and aboral ectoderm (Lapébie et al., 2014). It would also be interesting to test the importance of localized relaxation and contraction of the actomyosin cortex of different cell populations using optogenetics approaches (Krueger et al., 2018).

## Conclusions

Overall our study highlights how cell ingression and in particular the migration of ingressing cells relative to each other and to the blastocoel wall in *Clytia* not only provides the material to form the endodermal germ layer, but also generates forces that shape the embryo in the course of gastrulation. The polarized morphology of the canonical hydrozoan larva, including elongated shape and tapered posterior end, is thus tightly linked to the cell morphologenetic processes accompanying unipolar ingression, followed by migration and PCP-mediated interactions of the internalized cells.

## Acknowledgements

We thank all our research colleagues for their useful comments. We also thank Alexandre Jan and Laurent Gilletta for animal maintenance. Experimental work was supported by core CNRS and Sorbonne University funding to the LBDV. The Marine Resources Centre (CRBM) and PIV imaging platform of Institut de la Mer de Villefranche (IMEV) used in this study are supported by EMBRC- France (#ANR-10-INBS-02). We are very grateful to the Electron Microscopy Laboratory of the Shared Facilities Center of the MSU for great help and support. SEM was performed using the ‘Unique equipment setup 3D-EMС of Moscow State University’ supported by the Ministry of science and higher education of the Russian Federation, RFMEFI61919X0014. Y.K. research work was supported by federal project 0108-2019-0003 of the Koltzov Institute of Developmental Biology of the Russian Academy of Sciences.

